# Structural insights into agonist-binding and activation of the human complement C3a receptor

**DOI:** 10.1101/2023.02.09.527835

**Authors:** Manish K. Yadav, Ravi Yadav, Parishmita Sarma, Jagannath Maharana, Chahat Soni, Sayantan Saha, Vinay Singh, Manisankar Ganguly, Shirsha Saha, Htet A. Khant, Ramanuj Banerjee, Arun K. Shukla, Cornelius Gati

**Affiliations:** Department of Biological Sciences and Bioengineering, Indian Institute of Technology, Kanpur 208016, India; Bridge Institute, USC Michelson Center for Convergent Biosciences; University of Southern California, Los Angeles, CA, USA; USC Center of Excellence for Nano-Imaging, University of Southern California, Los Angeles, CA, USA; Department of Biological Sciences; University of Southern California, Los Angeles, CA, USA; Department of Chemistry, University of Southern California, Los Angeles, CA, USA

**Author notes:** Joint 1^st^ author (Manish K. Yadav, Ravi Yadav, Parishmita Sarma, Jagannath Maharana).

## Abstract

The complement cascade is an integral part of innate immunity, and it plays a crucial role in our body’s innate immune response including combating microbial infections. Activation of the complement cascade results in the generation of multiple peptide fragments, of which complement C3a and C5a are potent anaphylatoxins. The complement C3a binds and activates a G protein-coupled receptor (GPCR) known as C3aR while C5a activates two distinct receptors namely C5aR1 and C5aR2. Our current understanding of complement peptide recognition by their corresponding receptors is limited primarily to biochemical studies, and direct structural visualization of ligand-receptor complexes is still elusive. Here, we present structural snapshots of C3aR in complex with heterotrimeric G-proteins, with the receptor in ligand-free state, activated by full-length complement C3a, and a peptide agonist EP54, derived based on the carboxyl-terminal sequence of C5a. A comprehensive analysis of these structures uncovers the critical residues involved in C3a-C3aR interaction, and also provides molecular insights to rationally design carboxyl-terminal fragments of C3a and C5a to act as potent agonists of the receptor. Surprisingly, a comparison of C3a-C3aR structure with C5a-C5aR1 structure reveals diagonally opposite placement of these two complement peptides on their respective receptors, which helps explain the subtype selectivity of these complement peptides. Finally, taking lead from the structural insights, we also identify EP141, a peptide derived from the carboxyl-terminus of C3a, as a G-protein-biased agonist at C3aR. Taken together, our study illuminates the structural mechanism of complement C3a recognition by C3aR, and it also offers the first structural template for designing novel C3aR ligands with therapeutic potential for inflammatory disorders.

## Introduction

One of the key mechanisms through which the immune system combats pathogenic infections is the activation of the complement cascade^1–4^. It is an intricate network of plasma proteins including inflammatory peptides, proteases and integral membrane receptors that work in a concerted fashion^1–4^. The abnormal activation and levels of the complement components is also emerging as The complement fragments C3a and C5a generated from the proteolytic cleavage of complement C3 and C5, respectively, play a central role by priming and amplifying the immune response by recruiting the immune cells such as leukocytes and triggering the secretion of pro-inflammatory molecules, such as cytokines^1–4^. C3a and C5a bind to distinct seven transmembrane receptors (7TMRs) with C3a being selective for C3aR while C5a binds two different receptors namely, C5aR1 and C5aR2^5–8^. C3aR is a prototypical G protein-coupled receptor, which activates Gαi subtype of heterotrimeric G-proteins and recruits β-arrestins (βarrs) upon activation^9–12^ (Figure 1A). C3aR is expressed in multiple types of immune cells including mast cells, neutrophils and monocytes^13–16^. Increased levels of C3a have been implicated in multiple inflammatory disorders and C3a-induced C3aR signaling contributes to the pathophysiology of sepsis, arthritis, asthma and lupus^17–20^. Despite its fundamental importance, our current understanding of C3a-C3aR interaction and receptor activation is based primarily on biochemical studies^21,22^.

**Fig. 1.**
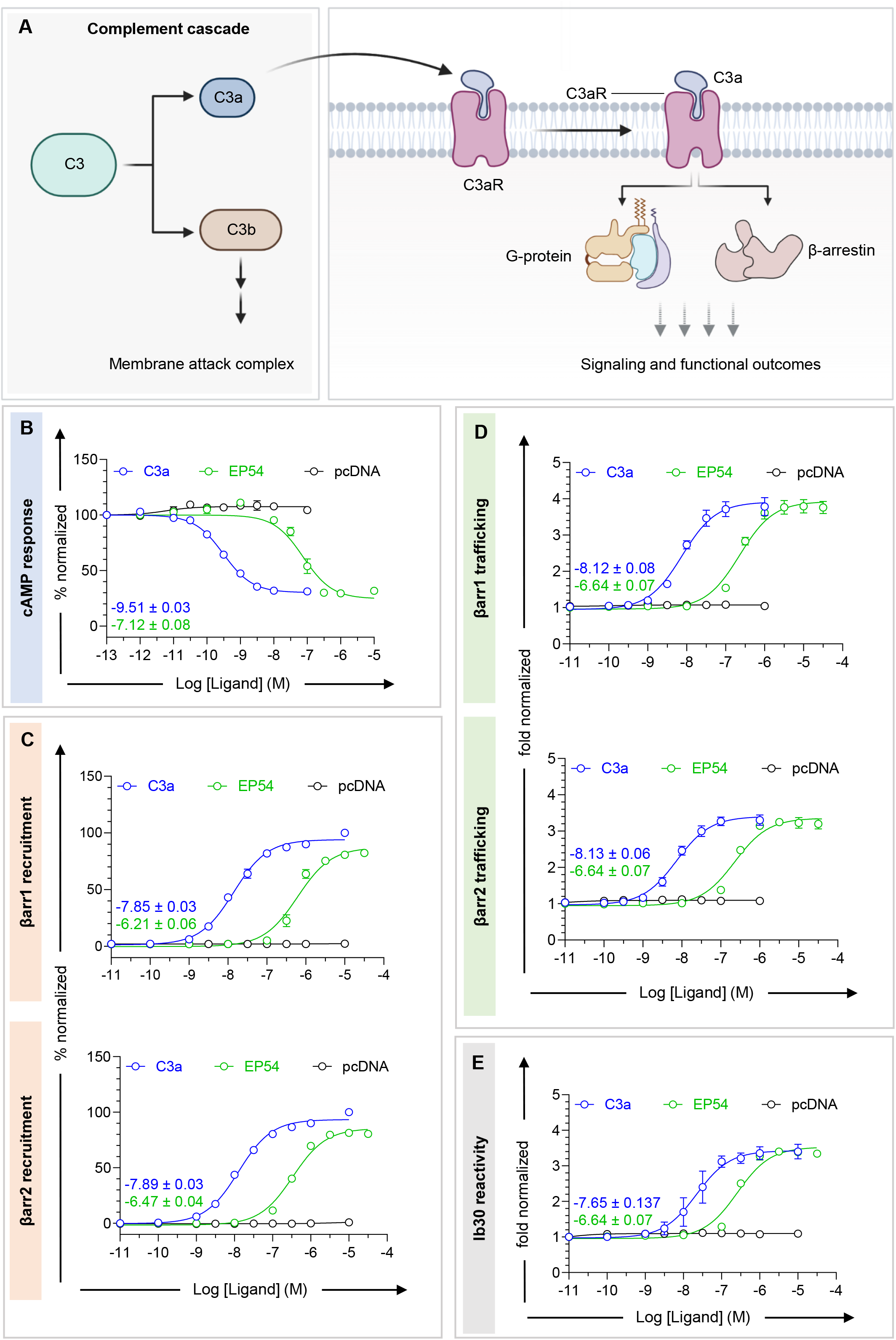
Overall C3aR signaling. **A.** Overview of the signaling cascade mediated by C3aR upon activation by C3a. **B.** To study Gαi-activation, forskolin-elevated decrease in cAMP level is measured using GloSensor assay downstream of C3aR in response to indicated ligands (mean±SEM; n=4; normalized with starting value for each ligand as 100%). **C.** βarr1/2 recruitment to C3aR in response to indicated ligands as measured by NanoBiT assay (Receptor-SmBiT+LgBiT-βarr1/2), respectively (mean±SEM; n=4; normalized with the luminescence signal at minimal ligand dose of each condition as 1). **D.** βarr1/2 trafficking to the endosomes downstream of C3aR in response to indicated ligands as measured by NanoBiT assay (Receptor+SmBiT-βarr1/2+LgBiT-FYVE) (mean±SEM; n=4; normalized with the luminescence signal at minimal ligand dose of each condition as 1). **E.** NanoBiT based assay used to assess Ib30 reactivity for βarr1 downstream of C3aR in response to indicated ligands (mean±SEM; n=4; normalized with the luminescence signal at minimal ligand dose of each condition as 1).

C3a is a 77 amino acid peptide with a four-helix architecture^23^ and previous studies have suggested a two-site binding mechanism to the receptor involving the 2^nd^ extracellular loop (ECL2) and the transmembrane core of the receptor^21,22^. Interestingly, peptides derived from, and modified based on, the carboxyl-terminus of both, C3a and C5a have been identified as full agonists of C3aR although their binding affinity and potency are significantly lower than C3a^24–28^. For example, EP54 and EP67, two decapeptides derived from the carboxyl-terminus of C5a, and several peptides derived from the carboxyl-terminus of C3a, exhibit full agonism at C3aR, in ERK1/2 MAP kinase phosphorylation assay^26^. These studies underscore the critical contribution of the carboxyl-terminus of C3a in eliciting transducer-coupling and downstream functional responses. However, direct structural visualization of agonist binding to C3aR still remains elusive, and a high-resolution structure of C3aR in inactive or active state have not been available thus far. This has limited our current structural understanding of agonist-induced C3aR activation and restricts the possibility of structure-guided design of novel therapeutics targeting this receptor.

In this backdrop, we present structural details of the human C3aR in complex with heterotrimeric G-proteins with the receptor in its ligand-free state, activated by the natural agonist C3a, and a small peptide agonist, EP54. These structures reveal molecular details of agonist-binding to C3aR, receptor activation, and G protein-coupling. Importantly, a comparative analysis with a C5a-C5aR1 structure elucidates strikingly distinct binding modalities between C3a and C5a on their respective receptors, and these structural insights also allow us to identify and rationalize G protein-bias exhibited by a C3a-derived peptide, EP141.

## Results and discussion

### Reconstitution of agonist-C3aR-G-protein complexes and overall structures

In order to elucidate the agonist-binding and activation of C3aR, we focused our efforts on its natural agonist C3a and the peptide agonist EP54. We first measured pharmacological profile of EP54 with C3a as a reference in G-protein and βarr assays (Figure 1B-E and Figure S1). We observed that EP54 behaved essentially as a full agonist for Gαi-coupling, as measured using cAMP response in GloSensor assay, and NanoBiT-based βarr1/2 recruitment and endosomal trafficking assay, albeit with lower potency (Figure 1B-D). Moreover, EP54 also induced an active conformation of βarr1 as measured using an intrabody sensor referred to as Ib30^29^(Figure 1E). To determine the structure of C3aR ternary complexes, we expressed and purified full-length human C3aR using the baculovirus expression system. Subsequently, we reconstituted C3a-C3aR-Gαoβ1γ2 and EP54-C3aR-Gαoβ1γ2 complexes stabilized by ScFv16^30^by combining purified components (Figure S2A-B). Negative-staining of these complexes suggested an overall uniform particle distribution with 2D class averages as expected for typical GPCR-G-protein assemblies (Figure S2D-E). We successfully determined the cryo-EM structures of these complexes at 3.19 Å and 2.88 Å, respectively (Figure 2 and S3-6). While we observed robust density of the ligand in the EP54-C3aR-Gαoβ1γ2 complex, surprisingly, we did not observe any discernible density for C3a in the case of C3a-C3aR-Gαoβ1γ2 complex (Figure S3-S4). Considering potential dissociation of C3a during purification and freezing steps as a plausible reason for the empty ligand binding pocket, we supplemented the reconstituted C3a-C3aR-Gαoβ1γ2 complex with additional C3a immediately before freezing samples for cryo-EM. This strategy allowed us to obtain a C3a-C3aR-Gαoβ1γ2 complex structure at 3.18 Å with a clearly discernible C3a density in the binding pocket (Figure S5). The coulombic maps of all the structures were sufficiently clear and allowed unambiguous placement of all components (Figure 2 and Figure S8). A comprehensive table outlining the resolved residues of these different structures is provided in Figure S9. The overall structures of C3aR reveal a canonical GPCR topology with the second extracellular loop (ECL2) adopting a β-hairpin conformation (Figure 2), and an RMSD of 0.35 Å between the C3a-bound and EP54-bound structures, suggesting a high degree of structural similarity (Figure S10A).

**Fig. 2.**
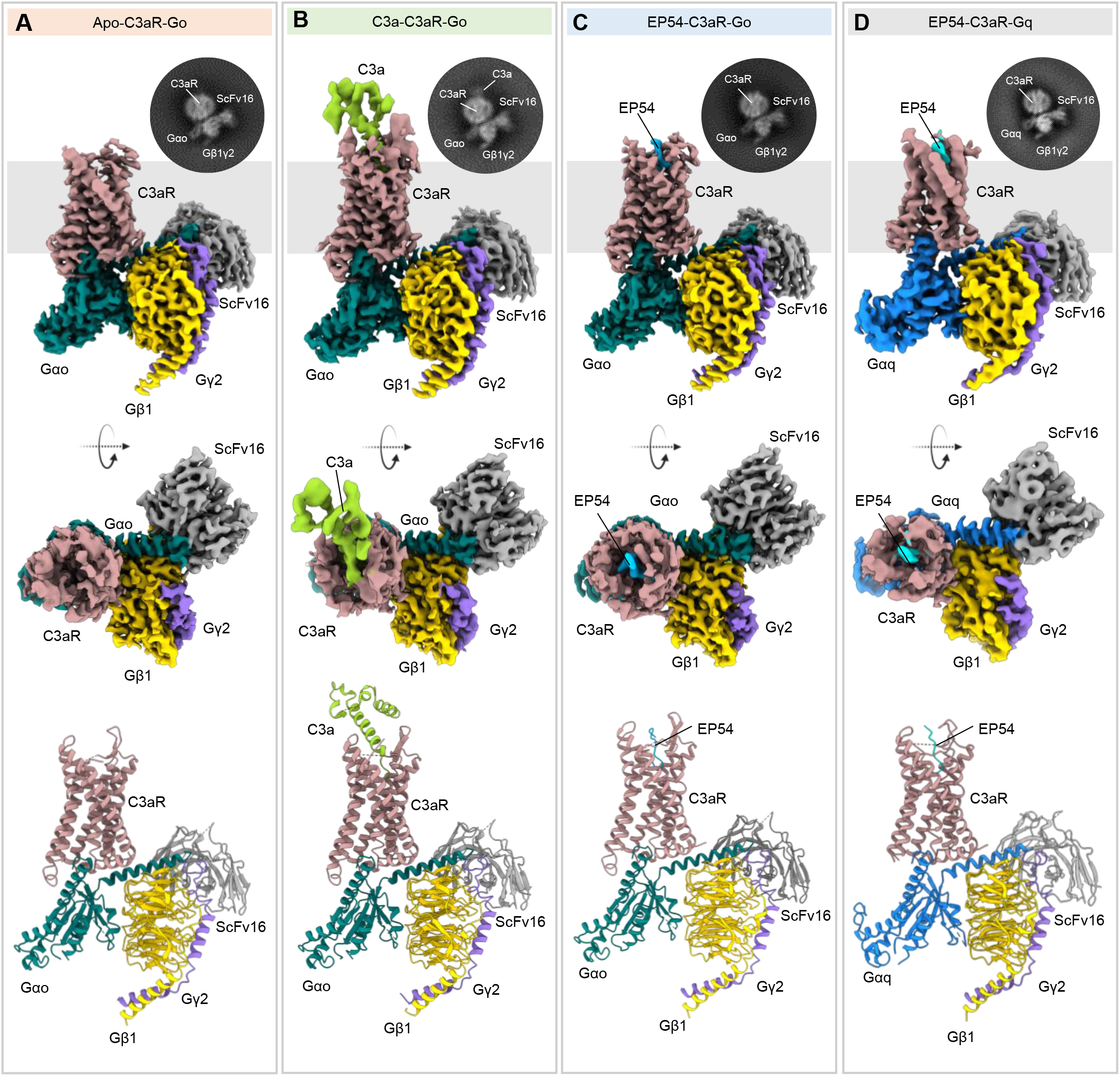
Structures of Apo-C3aR-Go, C3a-C3aR-Go and EP54-C3aR-Go complexes. **A, B, C, D.** Cryo-EM density maps of Apo-C3aR-Go (**A**), C3a-C3aR-Go (**B**), EP54-C3aR-Go (**C**), EP54-C3aR-Gq and **(D)** complexes along with its representative 2D class average in inset (top); the same electron density maps shown after rotating 90° (middle); Models of Apo-C3aR-Go, C3a-C3aR-Go, EP54-C3aR-Go and EP54-C3aR-Gq complexes shown in cartoon representation (bottom). Rosy brown, C3aR; teal, Gαo; yellow, Gαq; blue, Gβ1; purple, Gγ2; gray, ScFv16; green, C3a; cyan, EP54.

### Interaction of C3a and EP54 with C3aR

Complement C3a adopts a four-helix bundle structure stabilized by three disulfide bridges with the terminal arginine (Arg^77^) located at the end of helix 4 (H4)^23^ (Figure 3A). Site-directed mutagenesis studies coupled with ligand binding and second messenger assays have proposed a two-site model for C3a-C3aR interaction with the first site involving the 2^nd^ extracellular loop (ECL2) of the receptor, while the second site engages multiple residues from the extracellular side of the transmembrane helices^21,22^. It is noteworthy that although ECL2 of C3aR is extraordinarily long with more than 150 amino acids, functional studies have demonstrated that a large part of it is dispensable for C3a binding, as measured using calcium response as a readout^22^. In our C3a-C3aR-Gαoβ1γ2 structure, the stretch from Lys175^ECL2^ to Pro330^5.32^ is not resolved, likely due to inherent flexibility in this region and lack of further interactions with the receptor. In the structure, C3a retains the all-helical conformation in its basal state, and its carboxyl-terminus penetrates deep into the receptor core adopting a “hook-like” conformation with a buried surface area of 1,720 Å^2^ (Figure 3A-B).

**Fig. 3.**
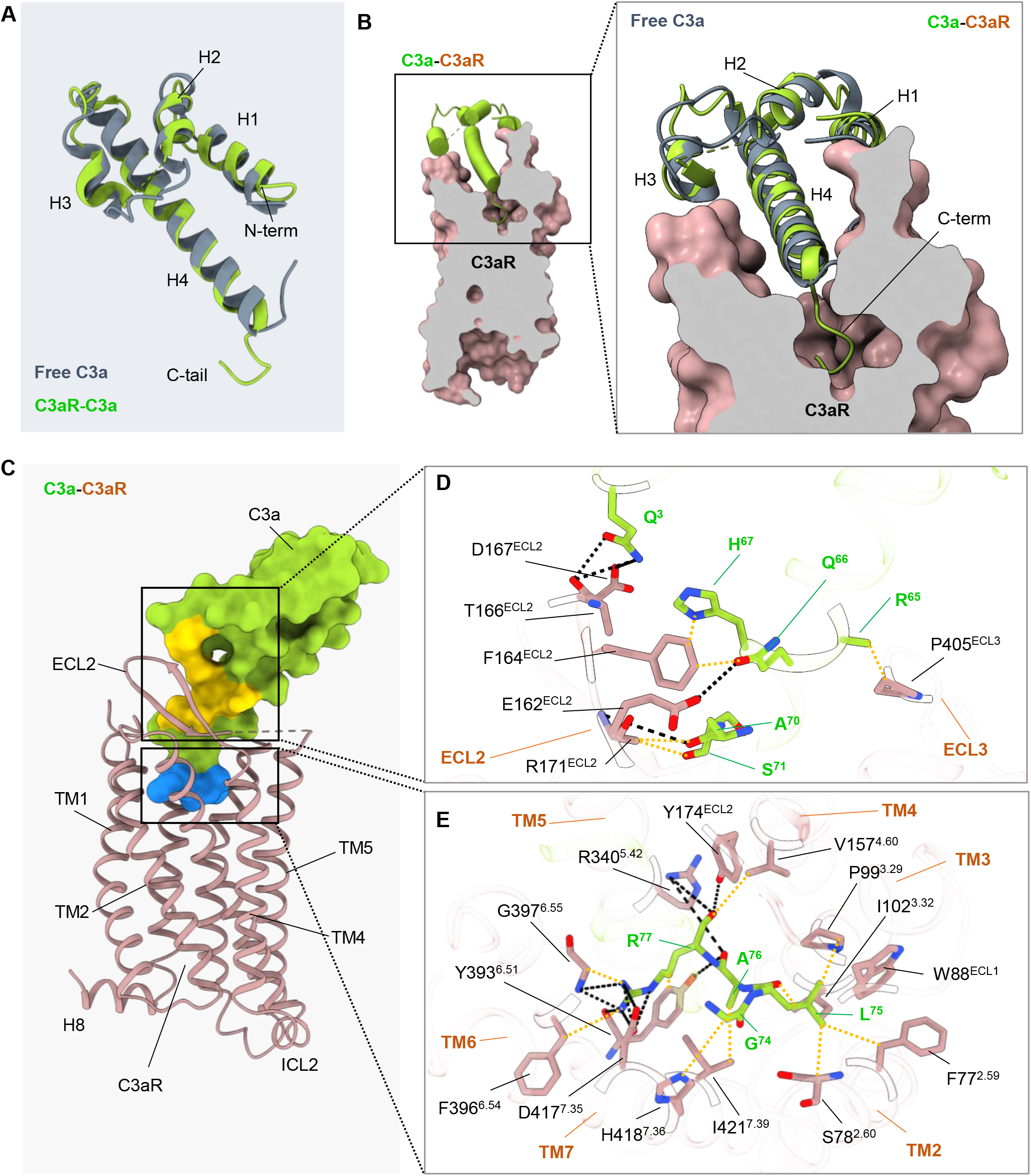
C3a binding to C3aR. **A.** Structure of C3a showing four helix bundle with a short C-terminal tail. Free C3a (PDB ID: 4HW5) has been superimposed on C3a bound to C3aR. **B.** The C-terminal tail of C3a adopts a hook like conformation upon entering deep into the orthosteric pocket of C3aR. C3aR shown in surface slice and C3a as cartoon representation. **C.** Side view of C3aR (cartoon) bound to its endogenous ligand C3a (surface). The binding sites have been assigned different colors: yellow (Site 1) and blue (Site 2). **D.** Details of interactions between ECLs of C3aR and globular domain of C3a at Site 1 (top), and Site 2 between TMs of C3aR and C-terminus of C3a (bottom) have been illustrated. Polar interactions have been depicted as black dotted lines, and non-bonded contacts in orange.

At the first site of C3a interaction, we observe that several residues from ECL2 of C3aR interact with His^67^-Ser^71^ stretch of H4 in C3a (Figure 3C-E). In addition, Gln^3^ in H1 of C3a forms ionic interactions with Thr166 and Asp167 in the ECL2 of C3aR, and it may contribute in stabilizing the ECL2 conformation (Figure 3D). Finally, Arg^65^ in H4 of C3a also forms a non-bonded contact with Pro405 in the ECL3 of C3aR (Figure 3D). At the second site, we expectedly observe an extensive engagement between the receptor residues in TM2, TM3, TM5, TM6 and TM7 with the L-G-L-A-R sequence in the carboxyl-terminus of C3a (Figure 3E). In particular, the terminal Arg^77^ of C3a localizes in a hydrophobic environment and engages with Tyr174^ECL2^ and Tyr393^6.51^ through hydrogen bonds, and with Asp417^7.35^ through a salt bridge. In addition, Arg^77^ also interacts with Val157^4.60^, Gly397^6.55^ and Ile421^7.39^ through van der Waals interactions, thereby enhancing the stability and positioning within the binding pocket (Figure 3E). The extensive interactions of Arg^77^ provide insights into its role in C3a engagement with C3aR. This is particularly revealing as a naturally-occurring proteolytic fragment of C3a (C3a^des-Arg^), devoid of terminal Arg^77^, almost completely loses the interaction with C3aR^31,32^, and our structural snapshot provides a molecular basis for this observation (Figure S11B-C). This is further supported by site-directed mutagenesis studies where Arg340Ala and Asp417Ala mutations in C3aR, which are critical interaction partners of Arg^77^ in C3a, diminish agonist-binding and receptor activation^21^.

Interestingly, we observe that the carboxyl-terminus of C3a in receptor-bound state exhibits a sharp turn starting at Ser^71^ compared to free C3a (PDB ID: 4HW5) that allows its penetration in the receptor core (Figure 3B). In addition to Arg^77^, C3a residues Gly^74^, Leu^75^ and Ala^76^ form hydrophobic contacts with Phe77^2.59^, Ser78^2.60^, Trp88^ECL1^, Pro99^3.29^, Ile102^3.32^, Ile421^7^.^39^ and His418^7.36^ of C3aR there are non-bonded contacts between Gly^74^-His418^7.36^, extensive hydrophobic contacts between Leu^75^ and Ser78^2.60^, Phe77^2.59^, Trp88^ECL1^, Pro99^3.29^ and Ile102^3^.^32^, and hydrophobic contact between Ala^76^ and Ile421^7.39^ (Figure 3E). These extensive interactions help stabilize and position the sharp kink in the carboxyl-terminus of C3a in the receptor core. This observation provides a structural rationale for the ability of peptides derived from the carboxyl-terminus of C3a to act as potent receptor agonists^25,33^, and therefore, also identifies ligand-receptor interactions that are critical and sufficient for driving downstream responses. A comprehensive detail of all the interactions between C3a and C3aR are listed in Figure S11A.

As mentioned earlier, peptides derived from, and modified based on, the carboxyl-terminal sequence of C3a and C5a act as potent agonists of C3aR^25,33^. The EP54-C3aR-Gαoβ1γ2 structure provides molecular insights into the interaction and efficacy of such peptides (Figure 4A). Similar to the carboxyl-terminus of C3a, EP54 also adopts a hook-like conformation and positions in an overall similar binding pocket on C3aR (Figure 4B-D). While there are a set of interactions that are common for C3a and EP54, we also observe multiple interactions that are specific to either C3a or EP54 (Figure 4E). The terminal arginine (Arg^10^) of EP54 exhibits extensive interactions with residues in several TMs of C3aR that are similar to those displayed by Arg^77^ in C3a (Figure 4B-C). For example, Arg^10^ in EP54 is localized in a negatively charged microenvironment surrounded by Tyr393^6.51^, Asp417^7.35^ and Cys420^7.38^, and forms hydrogen bond with Tyr174^ECL2^ of the receptor (Figure 4C). The stabilization of the ligand is further facilitated through an ionic interaction between side chain of Arg^10^ with Asp417^7.35^, a cation-π interaction with Tyr393^6.51^ and ionic bond formed by the carboxylate oxygen with the side chain of Arg340^5^.^42^ (Figure 4C). d-Ala^9^ of EP54 is engaged with the side chains of Ile102^3.32^, Met106^3^.^36^, Tyr393^6.51^ and Ile421^7^.^39^ through hydrophobic interactions, while Leu^8^ and Met^6^ forms hydrophobic contacts with His81^2.63^ and Ile102^3.32^ of C3aR, respectively. Interestingly, several C3a-interacting residues in ECL2 of C3aR also interact with EP54, suggesting at least a partial contribution of the first site in EP54 binding as well (Figure 4B). A comprehensive detail of all the interactions between EP54 and C3aR are listed in Figure S12.

**Fig. 4.**
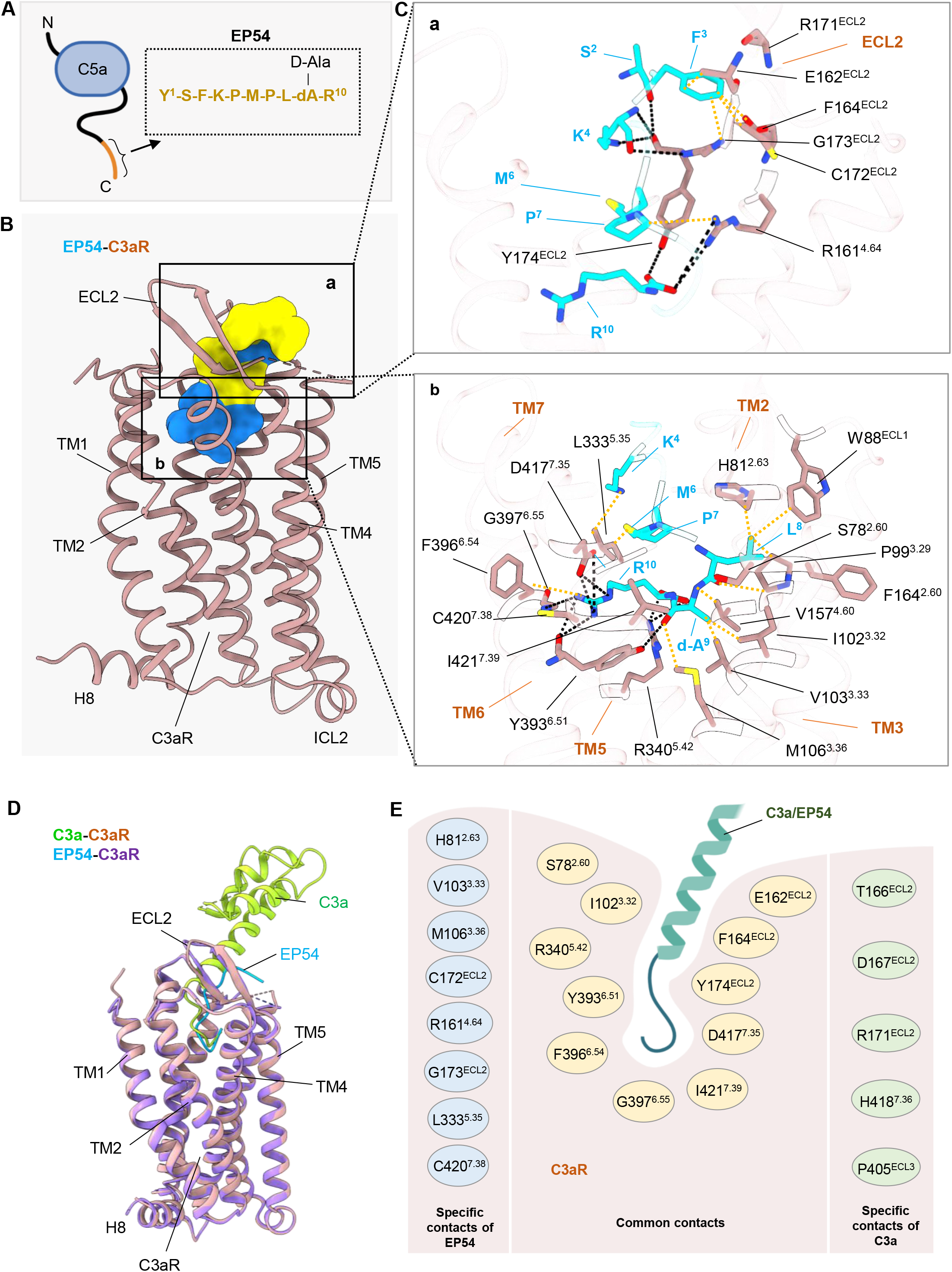
EP54 interactions with C3aR. **A.** Sequence of EP54 derived from the C-terminus of C5a. **B.** Side view of EP54 (surface) binding to C3aR (ribbon). Different sites of contact have been illustrated in different colors: yellow (Site 1), blue (Site 2). **C.** Details of interactions between ECLs of C3aR and EP54 at Site 1 (top), and at Site 2 between TMs of C3aR and EP54 (bottom). Polar interactions have been depicted as black dotted lines, and non-bonded contacts in orange. **D.** Structural superimposition of C3a/EP54 bound C3aR. **E.** Comparative analysis of common and specific interactions of C3a/EP54 with C3aR.

As mentioned above, we serendipitously determined the structure of ligand free C3aR-Gαoβ1γ2 complex, which exhibits an empty ligand binding pocket (Figure S13 and 14). Structural comparison between ligand-free and agonist-bound C3aR structures revealed differences in the overall positioning and conformation of some of the residues in the ligandbinding pocket. For example, Arg340^5.42^ of C3aR, which engages with Arg^77^ of C3a, displays a linear inward shift of about 5 Å, possibly due to a lack of constraints imposed by ligand binding (Figure S14B). Similarly, Arg161^4.64^ in TM4 of C3aR, which engages with Leu^8^ in EP54, exhibits a rotation of about 150° in the apo-C3aR to interact with the main chain of Lys96^3.26^ in TM3 (Figure S14C). Although several GPCRs exhibit constitutive activity, it remains to be determined whether this ligand-free-C3aR structure indeed represents a constitutively active conformation or it results simply due to the dissociation of C3a.

### Agonist-induced activation of C3aR and interface with G-proteins

The structures of C3aR in apo-, C3a- and EP54-bound states are overall similar, with an RMSD of <1 Å (Figure S13A). As a structure of C3aR in its inactive state is not available, we used the previously determined inactive state crystal structure of C5aR1^34,35^, which is a close phylogenetic homologue of C3aR, to identify activation-dependent structural features in C3aR. Superimposition of the C3aR structures determined here with that of antagonist-bound C5aR1 reveals anticipated conformational changes on the intracellular side of the receptor including an outward movement of TM6 by approximately 9 Å and an inward shift of TM7 by about 11 Å (Figure 5B). Interestingly, helix 8 in C3aR undergoes a dramatic movement of nearly 180° compared to the inactive C5aR1 structure, which potentially facilitates the opening of the transducer-coupling interface on the receptor (Figure 5B). In addition, the conserved microswitches in C3aR including DRY, NPxxY, CWxP and PIF motifs exhibit significant structural rearrangements upon activation as outlined in Figure 5C. Interestingly, these structural changes are similar in ligand free, EP54- and C3a-bound receptor, which could provide a molecular basis for constitutive activity observed in C3aR as well as full agonism of EP54 (Figure 5C). The overall interaction interface between the receptor and G protein is identical in all three structures (Figure 6A-C). The carboxyl-terminus of the α5 helix in Gαo adopts a loop conformation and inserts into the cytoplasmic core of the receptor with buried surface area of >2,200 Å^2^. The key interaction interface is composed of TM2, TM3, TM6, TM7, ICL2 and ICL3 in the receptor, and α5 helix, αN helix and αN–β1 loop of the Gαo subunit (Figure 6A-C and Figure S15-17). Specific interactions involve hydrogen bond formation between Asn347, Gly350, Cys351, Gly352, Leu353 and Tyr354 in the α5 helix of Gαo with Val123^3.53^, Asn57^2.39^, Arg120^3.50^, Thr375^6.33^ and Arg367^ICL3^ of the C3aR (Figure 6E-G). Additional interactions that contribute towards the receptor-G-protein interface include the interaction of residues from ICL3 with residues of α5 helix and α4-β6 loop of Gαo (Figure 6E-G). For example, Arg367 in ICL3 of C3aR forms hydrogen bonds with Tyr320 (β6 strand) and Tyr354 (α5 helix), polar interactions with Asn316 (α4-β6), and Asp341 (α5 helix) (Figure 6E-G) of Gαo.

**Fig. 5.**
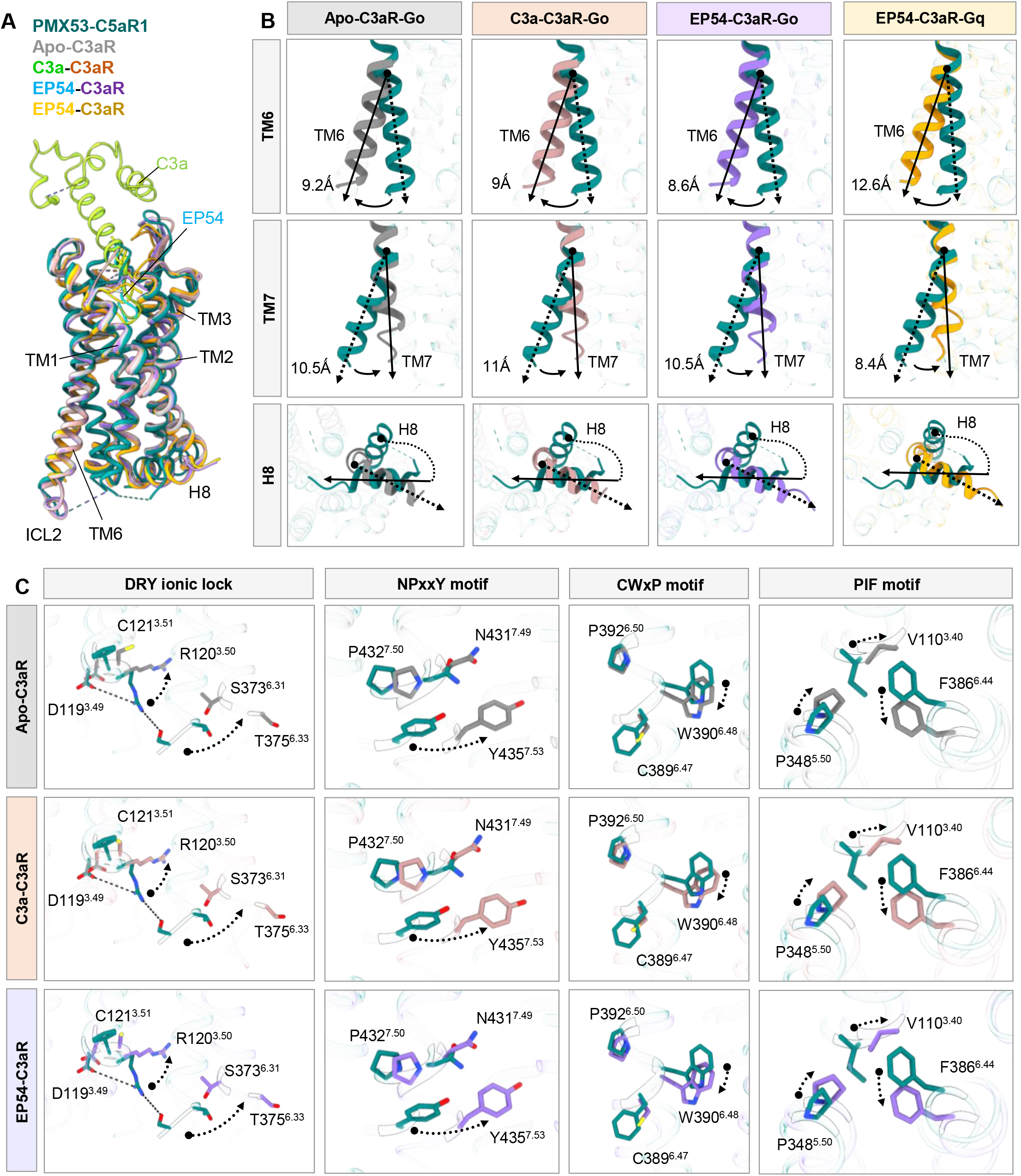
Conformation and activation hallmarks of C3aR. **A.** Structural superimposition of Apo and EP54/C3a bound C3aR with inactive C5aR1 (PDB ID: 6C1R). **B.** Dynamic changes in TMs of activated C3aR compared with the inactive state of C5aR1. **C.** Close-up views of the conserved DRY, NPxxY, CWxP and PIF motifs (left to right) showing conformational changes upon receptor activation. Polar contacts are depicted as black dashed lines.

### Structural insights into C3aR-Gαq coupling

In addition to Gαi-coupling, C3aR is also shown to couple to Gαq subtype of G-proteins^36^. Therefore, we also determined the structure of EP54-C3aR-Gαq complex at 3.57Å (Figure 2D) using the pipeline as described in Figure S18. The binding pocket for EP54 on the receptor, and its interaction with the receptor residues were also similar to that in the EP54-C3aR-Go complex with the major involvement of TM3-7, ECL2 and ECL3, and a buried surface area of 1,640Å^2^(Figure S19). For example, the terminal Arg^10^ of EP54 in the Gαq structure is positioned in a similar fashion as in EP54-C3aR-Gαo structure, and also aligns with the terminal Arg^77^ of C3a in C3aR-Gαo complex (Figure S19A-B). A complete list of all the interactions between EP54 and C3aR in the Gq complexes has been provided in Figure S20A. In addition, the overall structure of the receptor in this complex was similar to that in EP54-C3aR-Go complex with an RMSD of 0.753 Å for the Cα atoms (Figure S19C). At the receptor level, TM6 displays a slightly larger outward movement than in the Go complex (~12.5 Å vs. ~8.5 Å) while the inward movement of TM7 is a little smaller (~8.5 Å vs. ~10.5 Å) (Figure 5B). Compared to the inactive conformation of C5aR1, we also observe a large rotation of helix 8 in C3aR, and typical structure rearrangements in the conserved micro-switches as described earlier for Gαo-bound C3aR (Figure 5B-C). Finally, the overall engagement and interaction interface of C3aR and Gq is similar to that observed for C3aR-Go complex although there are some modest differences as well such as small outward displacement of α5 helix of Gαq (Figure 6D, H and S19C). Activation-dependent coupling of C3aR with Gq is facilitated primarily through the interactions involving TM2-TM4, TM6, and ICL2 on the receptor side, and the α5-helix from Gαq (Figure 6D, H). The key interactions at the receptor-Gq interface are presented in Figure 6H, and a comprehensive list of interacting residues is listed in Figure S20B.

**Fig. 6.**
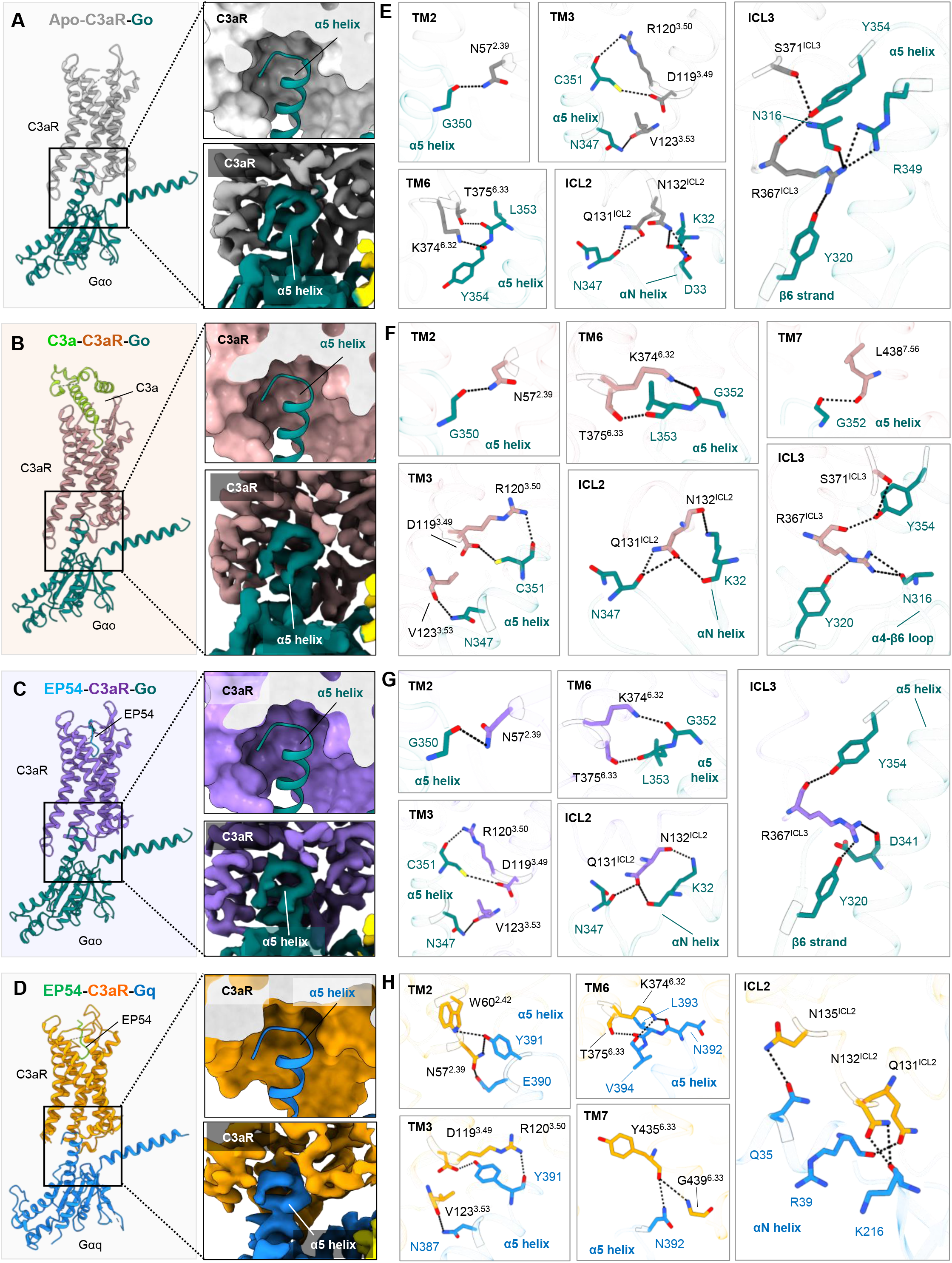
Interactions between C3aR and Gαo/Gαq. **A, B, C, D.** α5 helix of Gαo docks into the cytoplasmic core of C3aR. **E, F, G, H.** Magnified view of the interactions between TM2, TM3, TM6, TM7, ICL2 and ICL3 of C3aR with the Gαo/Gαq. Ionic bonds are depicted as black dashed lines.

### Distinct binding modalities of C3a and C5a on their respective receptors

We have recently determined the structure of C5a-C5aR1-Gαoβ1γ2 complex by cryo-EM^37^. In order to identify potential commonalities and differences between C3a and C5a binding to their respective receptors, we compared the overall structures and agonist-binding modes in C3a-C3aR and C5a-C5aR1. As presented in Figure 7A, the overall four-helix bundle in C3a and C5a remains unchanged upon receptor binding, however, there is a dramatic difference in the overall binding pose of C3a to C3aR and C5a to C5aR1 (Figure 7B-C and Figure S10C-D). The globular domain of C3a is tilted at an angle of 117° while that of C5a is tilted at an angle of 115° with respect to their respective carboxyl-terminus (Figure 7C). Moreover, the globular domains of C3a and C5a are oriented in opposite directions on the extracellular side of the receptor at an angle of 87° (Figure 7C). Still however, the carboxyl-terminus of both C3a and C5a adopt a hook-like conformation, share similar interacting residues, and position themselves in an analogous binding pocket in the receptor (Figure 7D-E). It is also noteworthy that while we observe a clear density for the N-terminus of C5aR1 and its interaction with C5a in the C5a-C5aR1 structure, we did not observe any discernible density for the N-terminus of C3aR in the structure. Taken together, the differential involvement of the N-terminus of the receptor and distinct orientation of C3a and C5a on C3aR and C5aR1 respectively, provides a structural basis of subtype selectivity between these two receptors. On the other hand, an overall similar interaction of the carboxyl-terminus of C3a and C5a with their respective receptors may help rationalize the cross-reactivity of peptides across these receptors^26^.

**Fig. 7.**
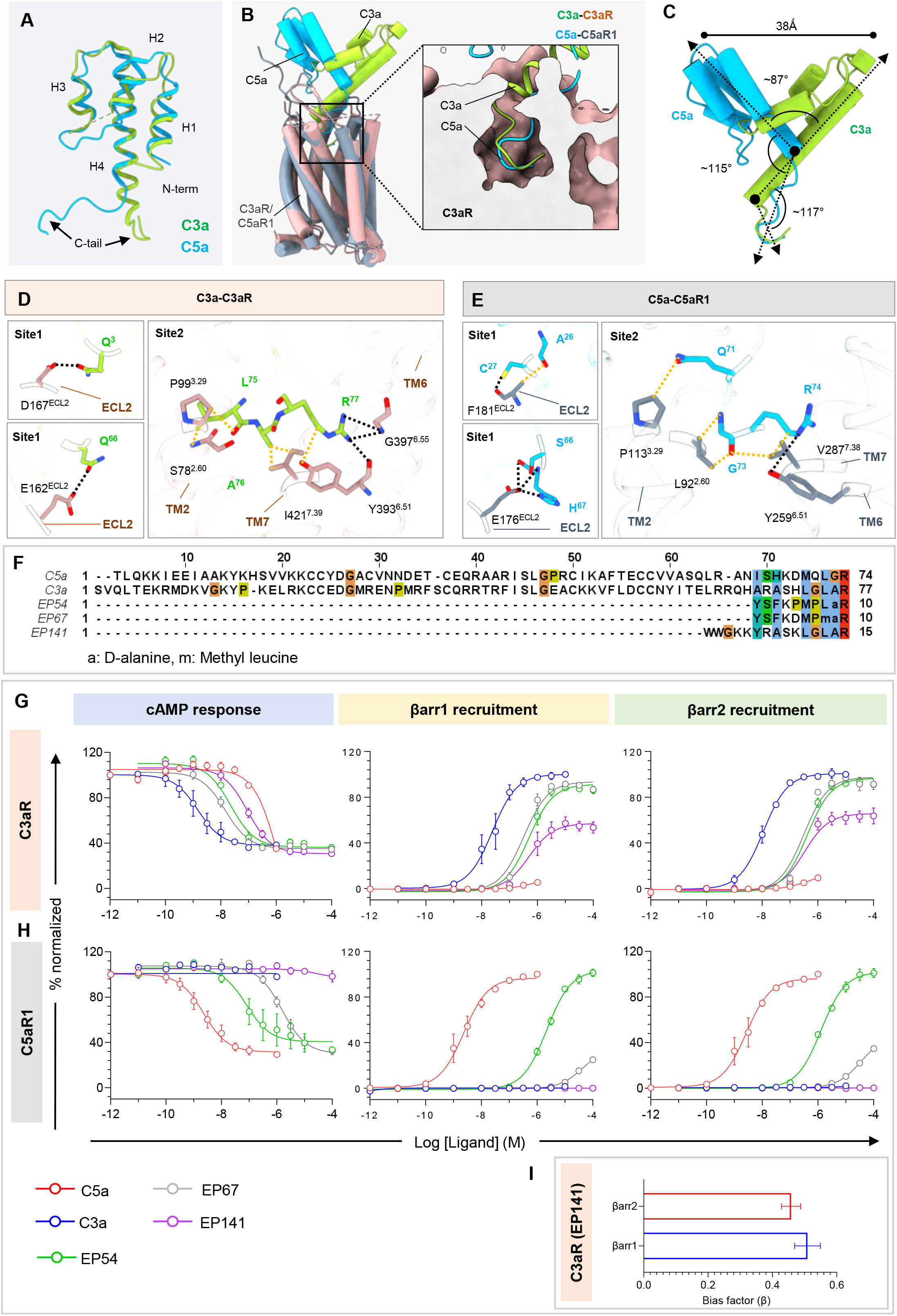
Cross-reactivity of C3a and C5a with complement receptors, C3aR and C5aR1. **A.** Structural superimposition of C3a and C5a shows conservation in the four helical bundle architecture with different conformation in the C-terminus upon binding to the receptor. **B.** Structural alignment of ligand bound C3aR and C5aR1 reveals conformational similarity in the C-terminus of their cognate endogenous ligands. C3aR in surface slice and ligands in cartoon representation. **C.** Differential positioning of C3a and C5a upon interaction with their respective receptors. **D, E.** Magnified view of the conserved contacts in the orthosteric pocket of C3aR and C5aR1. Ionic bonds are depicted as black dashed lines, and non-bonded contacts as orange. **F.** Sequence alignment of the complement peptides have been shown. The notations for unnatural, modified amino acids have been given below the alignment. **G.** G-protein activation and βarr1/2 recruitment were studied using GloSensor assay and NanoBiT-based assay (Receptor-SmBiT+LgBiT-βarr1/2) respectively, first panel: forskolin induced cAMP level downstream of C3aR in response to indicated ligands (mean±SEM; n=4; normalized with the lowest ligand concentration for each ligand as 100%), second panel: βarr1 recruitment to C3aR (mean±SEM; n=4) and third panel: βarr2 recruitment to C3aR (mean±SEM; n=3); normalized with the highest ligand concentration of C3a as 100%. **H.** G-protein activation and βarr1/2 recruitment downstream of C5aR1 in response to indicated ligands, first panel: forskolin induced cAMP level decrease downstream of C5aR1 in response to indicated ligands (mean±SEM; n=5; normalized with the lowest concentration of each ligand as 100%), second panel: βarr1 recruitment to C5aR1 (mean±SEM; n=5) and third panel: βarr2 recruitment to C5aR1 (mean±SEM; n=4); normalized with the highest ligand concentration of C5a as 100%. **I.** Bias factor (β value) determined taking C3a as reference elucidates the G-protein biased nature of EP141. Bias factor was calculated using https://biasedcalculator.shinyapps.io/calc/.

### Identification of a G-protein-biased peptide agonist at C3aR

Based on the structural insights wherein the terminal carboxyl-segment of C3a appears to be most critical for receptor binding and activation, we tested a set of peptide ligands including C3a and C5a on C3aR and C5aR1 in G-protein-coupling and βarr-recruitment assays (Figure 7F-H and Figure S21). Interestingly, while C3a did not exhibit any measurable transducer-coupling response at C5aR1, C5a exhibited some cross-reactivity with C3aR albeit only at high concentration in the G-protein-coupling assay (Figure 7F-H). More importantly, EP141 that is derived from, and modified based on, the carboxyl-terminus sequence of C3a showed full efficacy for C3aR in G-protein-coupling assay but only partial efficacy in βarr-recruitment (Figure 7F-H). In addition, while EP54 binds to both, C3aR and C5aR1, EP141 displayed robust selectivity for C3aR (Figure 7F-H). Taken together, these observations identify EP141 as a C3aR-selective, G-protein-biased agonist, and provide a framework for designing additional biased agonists at C3aR going forward (Figure 7I).

### Concluding remarks

The structural snapshots presented here elucidate the mechanistic basis of C3a recognition by its cognate GPCR, C3aR. In addition, we uncover the molecular mechanism that allows the carboxyl-terminal peptide fragments of C3a and C5a to bind and activate C3aR. Taken together with our accompanying manuscript^37^, we have visualized the structural basis of ligand-recognition and activation of two of the three complement receptors, which paves the way for structure-guided ligand discovery with subtype selective pharmacology and therapeutic potential in inflammatory conditions. Finally, similar to C5aR1, C3aR also exhibits efficient interaction with βarrs^9–11^, and therefore, it would be interesting to visualize C3a-C3aR-βarr complexes to better understand the details of transducer-coupling in the complement receptor system. It is also intriguing to note that unlike several GPCRs, βarrs play an inhibitory role in ERK1/2 MAP kinase activation for C3aR as reported earlier^10^. This makes C3aR a unique target to decipher conformational modalities in receptor-βarr complexes driving downstream signaling. Going forward, it would be interesting to visualize the third complement receptor, namely C5aR2, which couples exclusively to βarrs, without any detectable activation of G-proteins^38–40^.

## Data availability statement

Any additional information required to reanalyze the data reported in this paper is available from the corresponding author upon reasonable request.

## Authors’ contribution

MKY expressed and purified C3aR, and reconstituted the complex with G-proteins for negative-staining and cryo-EM; RY carried out cryo-EM screening, data collection and structure determination; PS generated all the constructs for functional assays and performed the cellular assays with some help from ShS; JM and RB carried out negative-staining, participated in structure determination, analyzed the structures and prepared the figures with help from MKY, PS and MG; CS started the expression, purification and reconstitution of C3aR complexes; VS and SS contributed in the purification of G-proteins, C3a and ScFv16; AKS and CG supervised and managed the overall project; all authors contributed to data analysis, interpretation and manuscript writing.

## Acknowledgements

This work is supported primarily by Science and Engineering Research Board (IPA/2020/000405 and SPR/2020/000408 and), an extramural grant from the Department of Biotechnology (DBT) (BT/PR29041/BRB/10/1697/2018) sanctioned under the Membrane Protein Structural Biology initiative, and National Bioscience Award (BT/HRD/NBA/39/06/2018-19). In addition, the research in A.K.S.’s laboratory is supported by the Senior Fellowship of the DBT Wellcome Trust India Alliance (IA/S/20/1/504916) awarded to A.K.S., Council of Scientific and Industrial Research [37(1730)/19/EMR-II], Indian Council of Medical research (F.NO.52/15/2020/BIO/BMS), Young Scientist Award from Lady Tata Memorial Trust, and IIT Kanpur. A.K.S. is EMBO Young Investigator. SS is supported by the Prime Minister’s Research Fellowship. We also thank Bhanupriya Panigrahi, Saloni Sharma and Gargi Mahajan for helping with protein purification. We thank T. Osinski at the USC Center for Advanced Research Computing (CARC) for support with computing resources. We acknowledge the Center of Excellence for Nano Imaging (CNI) at the University of Southern California for microscope time.

## Accession number

The cryo-EM maps and structures have been deposited in the EMDB and PDB with accession numbers EMD-35282 and PDB ID: 8I9S (Apo-C3aR-Go, Titan data), EMD-35259 and PDB ID: 8I97 (Apo-C3aR-Go, Glacios data), EMD-35275 and PDB ID: 8I9L (C3a-C3aR-Go, composite map), EMD-35257 and PDB ID: 8I95 (EP54-C3aR-Go), EMD-35263 and PDB ID: 8I9A (EP54-C3aR-Gq), respectively. The individual maps for composite map constitution have also been deposited to EMDB as EMD-35293 (C3aR-Go complex only, original map), and EMD-35294 (C3a only, original map).

## Materials and methods

### Methods

#### General reagents, plasmids, and cell culture

Unless otherwise stated, most standard reagents were purchased from Sigma Aldrich. Dulbecco’s Modified Eagle’s Medium (DMEM), Phosphate Buffer Saline (PBS), Trypsin-EDTA, Foetal-Bovine Serum (FBS), Hank’s Balanced Salt Solution (HBSS), and Penicillin-Streptomycin solution were purchased from Thermo Fisher Scientific. HEK293T cells (ATCC) were maintained in DMEM (Gibco, Cat. no: 12800-017) supplemented with 10% (v/v) FBS (Gibco, Cat. no: 10270-106) and 100 U ml^-1^ penicillin and 100 μg ml^-1^ streptomycin (Gibco, Cat. no: 15140122) at 37°C under 5% CO_2_. *Sf9* cells were maintained in protein-free cell culture media purchased from Expression Systems (Cat. no: 96-001-01) at 27°C with 135 rpm shaking. The cDNA coding region of C3aR was cloned in pcDNA3.1 vector with an N-terminal FLAG tag and in pVL1393 vector with an N-terminal FLAG tag followed by the N-terminal region of M4 receptor (residues 2-23). For the constructs used in NanoBiT assay, SmBiT fused at the C-terminus of the receptor were generated by sub-cloning in the lab, and other constructs have been described previously^29,40–45^. All DNA constructs were verified by sequencing from Macrogen. Recombinant human C3a and C5a (was purified from *Escherichia coli* following the previously described protocol with some modifications^23,46^. EP54, EP67, and EP141 were synthesized from GenScript.

#### GloSensor-based cAMP assay

G-protein activation on agonist stimulation was quantified by GloSensor assay using cAMP level as readout as described previously^47^. Briefly, HEK-293 cells were co-transfected with 5 μg of FLAG-tagged C3aR and C5aR1 along with 2 μg F22 plasmid. Post 16-18 hrs of transfection, cells were trypsinized and harvested, followed by resuspension in assay buffer composed of 1X HBSS, 20 mM of 4-(2-hydroxyethyl)-1-piperazineethanesulfonic acid (HEPES), pH 7.4, and D-luciferin (0.5 mg ml^-1^) (GoldBio, Cat. no: LUCNA-1G). Cells were seeded in 96-well flat bottom white plate (Corning) at a density of 200,000 cells per 100 μl and incubated at 37°C for 90 min followed by 30 min incubation at room temperature. Basal readings were taken before ligand stimulation. To study ligand-induced Gi activation, cells were treated with forskolin at 1 μM concentration before stimulation, and readings were recorded until maximum luminescence signal was obtained. For stimulation, ligand concentrations were prepared by serial dilution in 1X HBSS, 20 mM HEPES, pH 7.4. The cells were stimulated with indicated doses of respective ligands. Changes in luminescence were recorded using a microplate reader (BMG Labtech). Data were normalized by treating the minimum agonist concentration as 100% and plotted using nonlinear regression analysis in GraphPad Prism software.

#### Surface expression assay

To study receptor surface expression, whole cell-based receptor surface ELISA was performed as previously discussed^48^. Briefly, cells transfected with receptor construct for respective assays were seeded in a 24-well with 0.01% poly-D-Lysine pre-coated plate at a density of 0.2 million cells well^-1^ and incubated at 37°C for 24 hrs. After 24 hrs, the plate was removed, growth media was aspirated, and the plate was washed with ice-cold 1X TBS, followed by 20 min of fixation with 4% PFA (w/v in 1X TBS) on ice. After fixation, cells were washed thrice with 1X TBS (400 μl in each wash), followed by blocking with 1% BSA prepared in 1X TBS at room temperature for 90 min. Afterward, cells were incubated for 90 min with anti-FLAG M2-HRP (prepared in 1% BSA, 1:5000) (Sigma, Cat. no: A8592). Following antibody incubation, cells were washed thrice with 1% BSA (in 1X TBS). Thereafter, assay was developed by incubating cells with 200 μl of TMB-ELISA (Thermo Scientific, Cat no: 34028) until the light blue color appeared, followed by quenching with 100 μl of 1 M H_2_SO_4_ by transferring the blue-colored solution to a 96-well plate. Absorbance was measured at 450 nm using a multi-mode plate reader. Afterward, cells were washed twice with 200 μl of 1X TBS and then incubated with 0.2% Janus Green (Sigma; Cat no: 201677) w/v for 15 min. The excess stain was removed by three washes with distilled water. The stain was eluted by adding 800 μl of 0.5 N HCl per well. 200 μl of the eluted solution was transferred to a 96-well plate, and absorbance was recorded at 595 nm. The signal intensity was normalized by calculating the ratio of A450/A595 values. For data normalization, the ratio of A450/A595 values was calculated, followed by considering pcDNA transfected cells reading as 1, and receptor expression was calculated with respect to pcDNA. Data were analyzed in GraphPad Prism software.

#### NanoBiT-based βarr recruitment assay

Agonist-induced βarr1/2 recruitment to the plasma membrane downstream of C3aR was measured by luminescence-based enzyme-linked complement assay (NanoBiT-based assay) following the protocol described earlier^40,43,45^. Briefly, HEK-293 cells were transfected with C3aR (5 μg) harboring carboxyl-terminus fusion of SmBiT and βarr1/2 constructs (2 μg) with N-terminal fusion of LgBiT using transfection reagent polyethyleneimine (PEI) at DNA:PEI ratio of 1:3. Post 16-18 hrs of transfection, cells were trypsinized, harvested, and resuspended in the NanoBiT assay buffer (HBSS containing 0.01% BSA, 5 mM HEPES, pH 7.4, and 10 μM coelenterazine (GoldBio, Cat no: CZ05). After resuspension, 0.1 million cells per 100 μl were seeded in the flat-bottom white 96-well plate. The plate was incubated at 37°C for 90 min, followed by 30 min at room temperature. After 2 hrs of incubation, three cycles of luminescence reading were taken before ligand addition in a multi-plate reader (BMG Labtech). Ligand concentrations were prepared by serial dilution in 1X HBSS, 20 mM HEPES, pH 7.4. The cells were stimulated with indicated doses of respective ligands followed by measurement of luminescence signal using a multi-mode plate reader for 20 cycles, and average data of 5 cycles showing highest range of luminescence were used for analysis and presentation.

#### Agonist-induced endosomal trafficking of βarrs

Agonist-induced βarr1/2 trafficking to the endosomes was monitored using NanoBiT assay following the same protocol as described above for βarr recruitment. HEK-293 cells were transfected with βarr1/2 tagged with SmBiT at the N-terminus, and N-terminal LgBiT-tagged FYVE constructs were used for enzyme complementation. The amount of DNA for receptor βarr1/2 and FYVE was kept at 5 μg, 2 μg, and 5 μg, respectively.

#### Intrabody30 reactivity

Intrabody 30 (Ib30) reactivity towards receptor activated βarr1 was assessed by NanoBiT assay^29,40,44,45^. Cells were transfected with 5 μg C3aR, 2 μg SmBiT tagged βarr1, and 5 μg Ib30 tagged with LgBiT at the N-terminus. After 16-18 hrs of transfection, cells were seeded in 96 well plate at a density of 0.1 million well^-1^ and incubated at 37°C for 1 hr 30 min followed by 30 min at room temperature. The cells were then stimulated with varying doses of C3a and EP54 as shown in the graph).

#### Expression and purification of C3a and C3aR

Gene encoding hC3a and C5a were cloned in pET-32a(+) vector with a Trx-6X-His tag at the N-terminal end and purified following the previously described protocol for C5a purification with some modifications^23,46^. Briefly, freshly transformed *E. coli* shuffle cells were inoculated in 50 ml 2XYT media with 100 μg ml^-1^ampicillin for starter culture at 30°C. Overnight grown primary culture was inoculated into 1.5 L 2XYT media with 100 μg ml^-1^ ampicillin, and the culture was allowed to grow at 30°C. At O.D ~ 0.6, culture was induced with 1 mM IPTG and shifted to 16°C for overnight induction. Cells were harvested and incubated with 1 mg ml^-1^ lysozyme in 50 mM HEPES, pH 7.4, 300 mM NaCl, 20 mM Imidazole, 1 mM PMSF, and 2 mM Benzamidine) for two hrs at 4°C. Cells were disrupted with ultrasonication, and cell debris was removed with high-speed centrifugation. C3a/C5a was enriched on Ni-NTA resins using gravity flow. Nonspecific proteins were removed with extensive washing (50 mM HEPES, pH 7.4, 1 M NaCl, 20 mM Imidazole), and fusion-C3a/C5a was eluted with 50 mM HEPES, pH 7.4, 150 mM NaCl, 300 mM Imidazole. Trx-his tag was cleaved with 8-10 hrs TEV treatment (1:20 w/w, TEV: Fusion protein) at RT. Purified C3a and C5a were further cleaned using the Resource-S column and stored at −80°C with 10% final glycerol concentration.

Codon-optimized C3aR was expressed in *Spodoptera frugiperda (Sf9*) cells using the baculovirus expression system with an N-terminal FLAG tag to facilitate purification. The receptor was purified as described previously. Briefly, the insect cells were harvested at 72 hrs post-infection and lysed by sequential douncing in a low-salt buffer (20 mM HEPES, pH 7.4, 10 mM MgCl_2_, 20 mM KCl, 1 mM PMSF, and 2 mM Benzamidine), high salt buffer (20 mM HEPES, pH 7.4, 1 M NaCl, 10 mM MgCl_2_, 20 mM KCl, 1 mM PMSF, and 2 mM Benzamidine), and lysis buffer (20 mM HEPES, pH 7.4, 450 mM NaCl, 2 mM CaCl_2_, 1 mM PMSF, 2 mM Benzamidine and 2 mM Iodoacetamide). After lysis, the receptor was solubilized for 2 hrs at 4°C with continual stirring in a solution of 0.5% L-MNG (Anatrace, Cat. no: NG310) and 0.1% cholesteryl hemisuccinate (Sigma, Cat. no: C6512). Post solubilization, salt concentration was lowered to 150 mM, and the receptor was purified on M1-FLAG column. In order to remove nonspecific proteins from FLAG beads, three washes of low salt buffer (20 mM HEPES, pH 7.4, 2 mM CaCl_2_, 0.01% CHS, 0.01% L-MNG) were alternated with two washes of high salt buffer (20 mM HEPES, pH 7.4, 450 mM NaCl, 2 mM CaCl_2_, 0.01% L-MNG) after binding. The bound receptor was eluted with FLAG elution buffer (20 mM HEPES, pH 7.4, 150 mM NaCl, 0.01% MNG, 2 mM EDTA, and 250 μg ml^-1^ FLAG peptide) and alkylated with iodoacetamide to prevent aggregation. The purified receptor was concentrated using a 30 kDa MWCO concentrator and stored at −80 °C in 10% glycerol till further use. 100 nM of hC3a or 1 μM of EP54 were kept in all steps of receptor purification.

#### Expression and purification of G-proteins

Genes for miniGαo1 and miniGαq subunit were cloned in pET-15b(+) vector with an in-frame 6X-His tag at the N-terminal and expressed in *E. coli* BL21(DE3) cells^49,50^. A starter culture in LB media was grown at 37 °C for 6-8 hrs at 220 rpm. This was followed by an overnight primary culture at 30°C with 0.2% glucose supplementation.15 ml primary culture was inoculated in 1.5 L TB (Terrific Broth) media and induced with 50 μM IPTG at an O.D of 0.8 and cultured at 25°C for 18-20 hrs. Cells were lysed in lysis buffer (40 mM HEPES, pH 7.4, 100 mM NaCl, 10 mM Imidazole, 10% Glycerol, 5 mM MgCl_2_, 1 mM PMSF, 2 mM Benzamidine) in the presence of 1 mg ml^-1^ lysozyme, 50 μM GDP and 100 μM DTT. Cell debris was pelleted down by centrifuging at 18000 rpm for 30 min at 4°C. Protein was enriched on Ni-NTA bead and after washing extensively with wash buffer (20 mM HEPES, pH 7.4, 500 mM NaCl, 40 mM Imidazole, 10% Glycerol, 50 μM GDP and 1 mM MgCl_2_), eluted with elution buffer (20 mM HEPES, pH 7.4, 100 mM NaCl, 10% Glycerol, 500 mM Imidazole). Eluted protein was pooled and stored at −80°C in 10% glycerol till further use. The gene encoding the Gβ1 subunit with an in-frame C terminal 6X-His tag and the Gγ2 subunit was expressed in *Sf9* cells using the baculovirus expression system. Cells were harvested 72 hrs after infection and resuspended in lysis buffer (20 mM TrisCl, pH 8.0, 150 mM NaCl, 10% Glycerol, 1 mM PMSF, 2 mM Benzamidine and 1 mM MgCl_2_). The cells were lysed via douncing and centrifuged at 4°C for 40 min at 18000 rpm. Pellet was resuspended and dounced in solubilization buffer (20 mM Tris-Cl, pH 8.0, 150 mM NaCl, 10% Glycerol, 1% DDM, 5 mM β-ME, 10 mM Imidazole, 1 mM PMSF, and 2 mM Benzamidine) and solubilized at 4°C under constant stirring for 2 hrs. Cell debris was pelleted down by centrifuging at 20000 rpm for 60 min at 4 °C. Protein was enriched on Ni-NTA resin, and after extensive washing with wash buffer (20 mM Tris-Cl, pH 8.0, 150 mM NaCl, 30 mM Imidazole, 0.02% DDM), the protein was eluted with 300 mM Imidazole in 20 mM Tris-Cl, pH 8.0, 150 mM NaCl, 0.01% MNG. Eluted protein was concentrated with a 10 kDa MWCO concentrator (Cytiva Cat no: GE28-9322-96) and stored at −80°C with 10% glycerol.

#### Expression and purification of ScFv16

Gene encoding ScFv16 was cloned in pET-42a(+) vector with an in-frame N-terminal 10X-His-MBP tag followed by a TEV cleavage site and expressed in *E. coli* Rosetta (DE3) strain^30^. Overnight primary culture was sub-cultured in 1L 2XYT media supplemented with 0.5% glucose and 5 mM MgSO_4_. At O.D_600_ ~0.6, culture was induced with 250 μM IPTG for 16-18 hrs at 18°C. Cells were resuspended in 20 mM HEPES, pH 7.4, 200 mM NaCl, 2 mM Benzamidine, and 1 mM PMSF and incubated at 4°C for 1 hr with constant stirring. Cells were disrupted by ultrasonication, and cell debris was removed by centrifugation at 18000 rpm for 40 min at 4°C. Protein was enriched on Ni-NTA resins, and non-specifically bound proteins were removed by extensive washing (20 mM HEPES, pH 7.4, 200 mM NaCl, 50 mM Imidazole). Bound protein was eluted with 300 mM Imidazole in 20 mM HEPES, pH 7.4, 200 mM NaCl. Subsequently, Ni-NTA elute was enriched on amylose resin (NEB, Cat. no: E8021L) and washed with buffer (20 mM HEPES pH 7.4, 200 mM NaCl) to remove nonspecific proteins. Protein was eluted with 10 mM maltose (prepared in 20 mM HEPES, pH 7.4, 200 mM NaCl), and the His-MBP tag was removed by overnight treatment with TEV protease. Tag-free ScFv16 was recovered by passing TEV-cleaved protein through Ni-NTA resin. Eluted protein was concentrated and cleaned by size exclusion chromatography on Hi-Load Superdex 200 PG 16/600 column (Cytiva Life sciences, cat. no: 17517501). Purified protein was flash-frozen and stored at −80°C with 10% glycerol.

#### Reconstitution of the C3a/EP54-C3aR-Gαo/Gαq-Gβγ-ScFv16 complexes

Purified C3aR was incubated with 1.2 molar excess of Gαo1, Gβ1γ2, and ScFv16 at room temperature for two hours in the presence of 25 mU ml^-1^ apyrase (NEB, cat. no: M0398S) and either C3a or EP54 for complex formation. The G-protein complex was separated from unbound components by loading on Superose 6 increase 10/300 GL SEC column and analyzed by SDS page. Complex fractions were pooled and concentrated to ~10 mg ml^-1^ using a 100 MWCO concentrator (Cytiva, Cat. no: GE28-9323-19) and stored at −80°C until further use.

#### Single particle, negative-stain EM

In order to confirm homogeneity and complex formation, negative stain electron microscopy was performed on all the samples before proceeding on with grid preparation for cryo-EM data collection. The individual samples were diluted to 0.02 mg/ml just prior to grid preparation and 3 μl of the samples were dispensed onto the carbon side of a glow discharged carbon/formvar coated 300 mesh Cu grids (PELCO, *Ted Pella*). The extra protein sample was blotted off after incubation for 1 min using a filter paper. The grid with the adsorbed protein sample was then touched on a first drop of 0.75% uranyl formate stain, and immediately blotted off using a filter paper. The grid was then touched onto a second drop of stain and moved gently in a rotating fashion for 30 s to increase the efficiency of staining. The grid so prepared was then left in a desiccator or on the bench in a petri-plate for air drying. Data collection was performed with a FEI Tecnai G2 12 Twin TEM (LaB6) operating at 120 kV and equipped with a Gatan CCD camera (4k x 4k) at 30,000x magnification. Processing of the collected dataset was performed with Relion 3.1.2^51^ where almost 10,000 particles were autopicked and subjected to reference free 2D classification, generating the 2D class averages.

#### Cryo-EM grid preparation and data collection

Purified ligand-C3aR-Gαo complexes were applied onto glow-charged grids (EasiGlow, 20 mA current with 40 s glow and 10 s hold) at ~10 mg ml-1 concentration and blotted for 5-7 s followed by plunge-freezing into liquid ethane using a Vitrobot MarkIV (Thermo Fisher Scientific, USA) operating at 100 % humidity. Briefly, ligand free-C3aR-Gαo and EP54-C3aR-Gαo complexes were applied onto Quantifoil R1.2/1.3 Au 300-mesh grids and blotted for 5 s at 4°C. For C3a-C3aR-Go and EP54-C3aR-Gq complexes, the sample was applied onto Quantifoil R1.2/1.3 Au 200-mesh grids and blotted for 5-7 s at either 4 or 22°C. Cryo-EM data were collected using a Glacios microscope operating at 200 kV with a Falcon 4 direct electron detector operating in counting mode at nominal magnification of 150,000 resulting in pixel size of 0.92 Å using EPU. An additional dataset of ligand free-C3aR-Go was collected using Titan Krios operating at 300 kV with a Gatan K3 Summit direct electron detector operating in counting mode at 105,000 magnifications with a pixel size of 0.86 Å (2-fold binned) using SerialEM. Movies were recorded with a defocus range of −0.8 to −3.0 μm and total dose of ~50 e-/A2. Additional data collection parameters are listed in Figure S7. Total 4,614 movies for EP54-C3aR-Go, 4990 movies for EP54-C3aR-Gq, 20,051 movies for C3a-C3aR-Go 4,611 movies for ligand-free-C3aR-Go (Glacios) and 5,740 for ligand free-C3aR-Go (Titan) were recorded.

#### Image processing and map construction

The overall cryo-EM data processing pipeline for EP54-C3aR-Gαo, C3a-C3aR-Gαo, Apo-C3aR-Gαo and EP54-C3aR-Gαq are shown in Figure S3-6, S18. Data processing steps were performed in cryoSPARC version 4.0^52^ unless otherwise stated. Cryo-EM movie stacks were aligned using Patch motion correction (multi) followed by CTF estimation with Patch CTF estimation (multi). Micrographs were curated based on CTF resolution and selected micrographs were used for particle picking using blob-picker. The picked particles were extracted (box size of 416 px and fourier cropped to 64 px) followed by iterative 2D classification. 2D classes with secondary structure features were selected and the particles were re-extracted. A subset of particles was used for multi-class ab-initio reconstruction. The clean particle stack was subjected to iterative heterogeneous refinement using ab-initio models as reference. The particles/maps from heterogenous refined were further subjected to non-uniform and subsequently to local refinement with appropriate mask.

For the EP54-C3aR-Gαo complex dataset, a clean particle stack of 767,052 from the 2D classification step was selected and re-extracted with a box size of 360 px and fourier cropped to 256 px (pixel size of 1.495 Å). This particle stack was subjected to ab-initio reconstruction, followed by heterogenous refinement with C1 symmetry. Total 600,173 high-quality particles with evident structural features were selected and subjected to non-uniform refinement with C1 symmetry, followed by local refinement with a mask on the micelle. This led to a reconstruction with a global estimated resolution of 2.88 Å (voxel size of 0.92 Å) as determined by gold standard Fourier Shell Correlation (FSC) using the 0.143 criterion. Local resolution estimation was performed with the Blocres sub-program within cryoSPARC version 4.0. Maps were sharpened using the “Autosharpen” sub-program within the Phenix suite^53^ for better visualization and model building.

For the C3a-C3aR-Gαo dataset, post 2D classification, 922,698 selected particles were extracted with a box size of 416 px and fourier cropped to 360 px (pixel size of 1.063 Å). This clean particle stack was then subjected to ab-initio reconstruction and heterogeneous refinement. The 3D class containing 418,953 particles with evident secondary features were subjected to non-uniform refinement, followed by local refinement with mask to exclude the micelle resulted in a final map at 3.18 Å resolution (voxel size of 1.063 Å) according to the gold standard FSC criterion of 0.143. The density corresponding to C3a was noisy, therefore, local refinement was performed with a mask covering the C3a to improve the interpretability of the map in this region yielding a map with global resolution of 4.55 Å. The two local refinement maps were combined to obtain a composite map which was used for subsequent model building.

For the apo-C3aR-Gαo complex, two independent datasets were collected – one with Titan Krios microscope operating at 300 kV and the other with a Glacios microscope operating at 200 kV. For the Glacios dataset, 2D class averages with clear secondary features were selected to prepare a sub-set of 693,550 particles for further processing. Subsequent ab-initio reconstruction and 3D/Heterogeneous classification with C1 symmetry yielded the best class with 464,408 particles which was further refined to an overall resolution of 3.19 Å (voxel size of 1.063 Å) with NU refinement according to the gold standard FSC criterion of 0.143. For the apo-C3aR-Gαo Titan dataset, total 754,251 particles corresponding to 2D averages with clear secondary structural features were selected, re-extracted with a box size of 416 px and Fourier cropped to 256 px (pixel size 1.40 Å). This clean particle stack was used for multiclass ab-initio reconstruction followed by heterogeneous classification. Non-uniform refinement with 327,193 particles from the best 3D class yielded a map with a final estimated global resolution of 3.26 Å (voxel size of 1.40 Å).

Initial processing of EP54-C3aR-Gq micrographs showed limited top/bottom views. Therefore, a conventional neural network-based method TOPAZ^54^, implemented in cryoSPARC, was used for particle picking. Total 2,007,547 particles were picked, extracted and subjected to iterative rounds of reference-free 2D classification followed by multiclass ab-initio reconstruction/heterogeneous refinement. The best 3D class with 101,400 particles was selected for subsequent non-uniform refinement, which yielded a map of 3.57 Å resolution based on the gold standard FSCat0.143 criterion.

#### Model building and refinement

Coordinates of C5aR1 receptor, Gαo, Gβ1 and Gγ2 from the cryo-EM structure of C5a-pep-C5aR1-Gαo (PDB ID: 8HPT) were used as initial models to dock into the EM density of EP54-C3aR-Gαo complex using the “Fit in map” extension in Chimera^55,56^. The fitted model along with the map was imported into COOT^57–59^ for manual rebuilding of the model along with the ligand (EP54), with iterative rounds of real space refinement in Phenix^53^. This yielded a final refined model with 94.95 % of the residues in the most favoured region and 5.05 % in the allowed region of the Ramachandran plot. For the C3a-C3aR-Gαo complex map, coordinates of the receptor, Gαo, Gβ1 and Gγ2 from EP54-C5aR1-Gαo complex were used as initial models. The coordinates of C3a were obtained from a previously solved crystal structure of C3a (PDB ID: 4HW5). Chimera was used to dock the individual components in the cryo-EM map and obtain a merged model. The combined model was manually rebuilt in COOT^57–59^ and subjected to multiple rounds of real space refinement in Phenix. The final refined model showed good validation statistics with 95.44 % in the favoured region and 4.56 % in the allowed region of the Ramachandran plot. Similarly, for EP54-C3aR-Gq and both the ligand free-C3aR-Gαo cryo-EM maps, the coordinates corresponding to the individual components were obtained from the model of EP54-C3aR-Gαo which were used to fit into the coulombic map with the “Fit in map” extension in Chimera. Iterative rounds of manual adjustment and building in COOT^57–59^ and refinement with real space refine in Phenix resulted in the final model with excellent validation statistics with no Ramachandran outliers. For ligand free-C3aR-Gαo, the model obtained from the Glacios data was used for analysis of the ligand free structure due to its higher resolution and more interpretable map. Data collection and refinement statistics have been included in the supplemental information. Chimera or ChimeraX software^55,60^ was used to prepare all figures used in the manuscript. Buried surface area has been calculated with the PDBePISA webserver^61^.

