## Supplemental Figures for "Structural insights into agonist-binding and activation of the human complement C3a receptor"

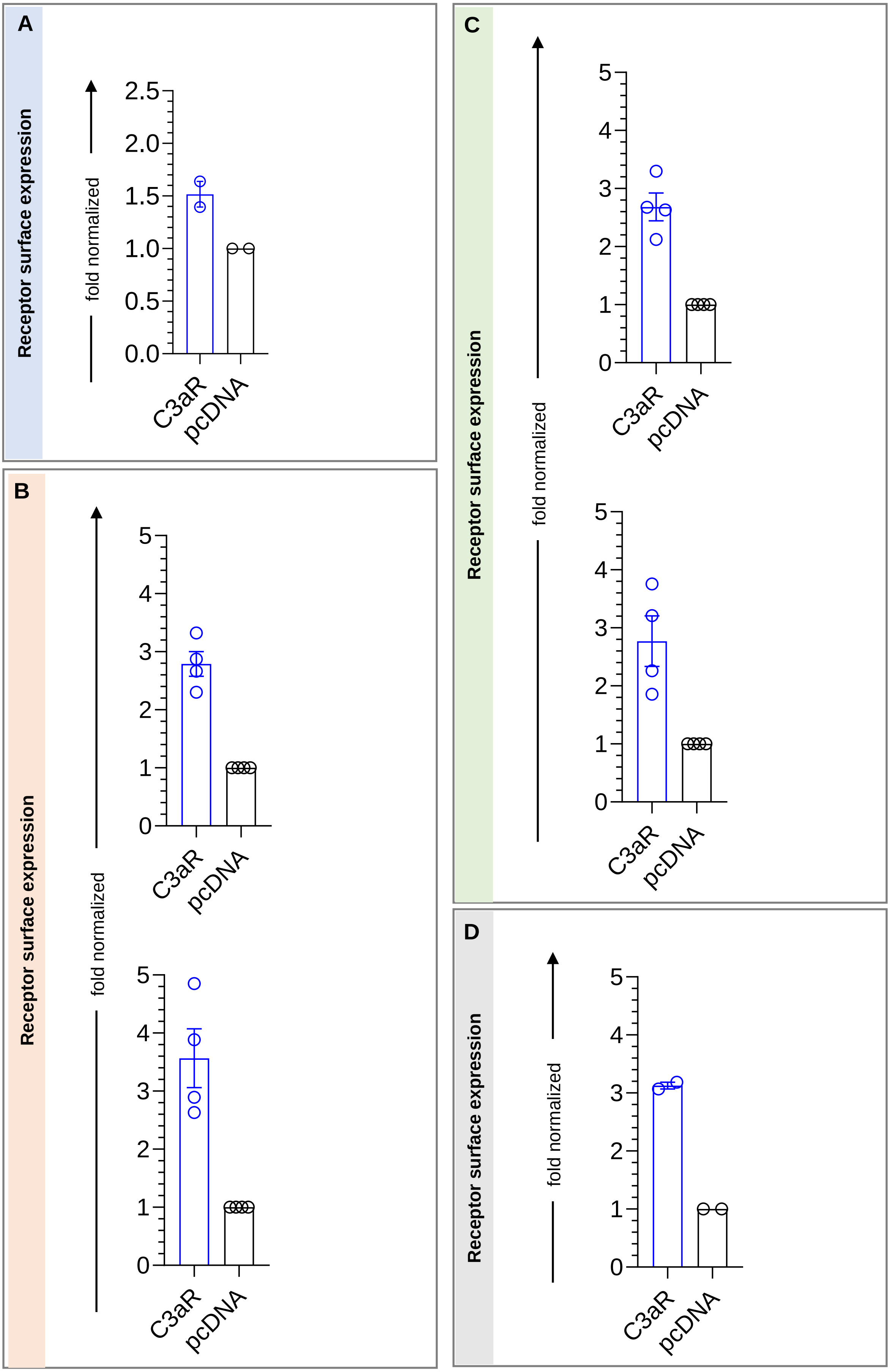
A
Receptor surface expression
fold normalized
2.5
2.0
1.5
1.0
0.5
0.0
C3aR
pcDNA
B
Receptor surface expression
fold normalized
5
4
3
2
1
0
C3aR
pcDNA
C
Receptor surface expression
fold normalized
D
Receptor surface expression
fold normalized

**Figure S1: Surface expression of C3aR in different assays.**

**A.** Surface expression of C3aR in GloSensor measured using whole cell ELISA (mean±SEM; n=2; normalized as fold over mock-transfection). **B.** Surface expression of C3aR in βarr1/2 recruitment assay (mean±SEM; n=4; normalized as fold over mock-transfection). **C.** Surface expression of C3aR in βarr1/2 endosomal trafficking assay (mean±SEM; n=4; normalized as fold over mock-transfection). **D.** C3aR surface expression in Ib30 reactivity NanoBiT assay (mean±SEM; n=2; normalized as fold over mock-transfection).

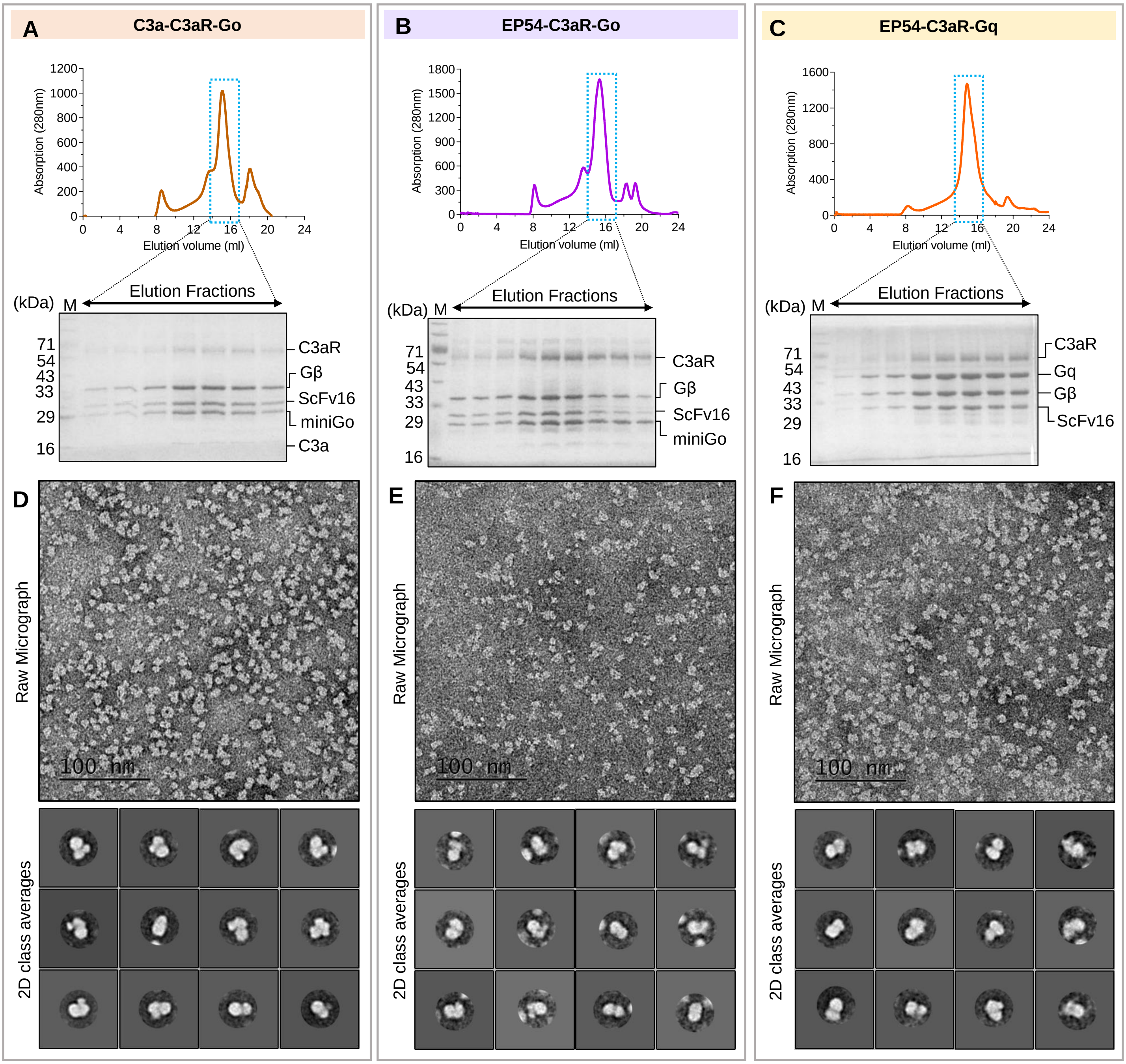
A
C3a-C3aR-Go
B
EP54-C3aR-Go
C
EP54-C3aR-Gq
Absorption (280nm)
Elution volume (ml)
Elution Fractions
(kDa)
M
71
54
43
33
29
16
C3aR
Gβ
ScFv16
miniGo
C3a
Gq
D
E
F
Raw Micrograph
100 nm
2D class averages

**Figure S2: Purification of EP54 and C3a bound C3aR-Go/Gq complexes and visualization through negative staining EM.**

**A, B, C.** Size Exclusion Chromatography profile and SDS-PAGE analysis of C3a-C3aR-Go, EP54-C3aR-Go and EP54-C3aR-Gq complex respectively. **D, E, F.** Negatively stained raw micrographs of C3a-C3aR-Go, EP54-C3aR-Go and EP54-C3aR-Gq complex respectively.

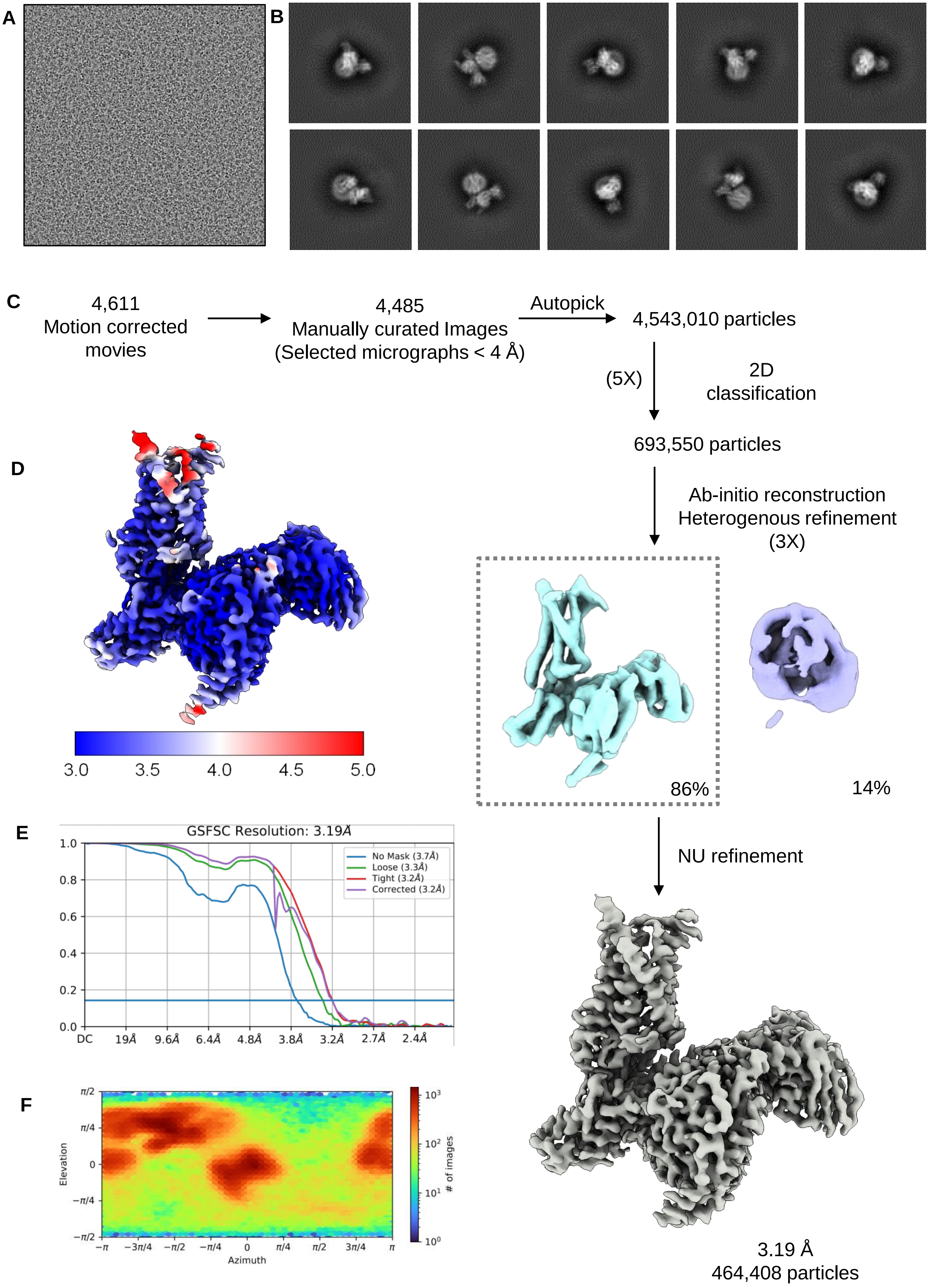
A
B
C
4,611
Motion corrected
movies
4,485
Manually curated Images
(Selected micrographs < 4 Å)
Autopick
4,543,010 particles
(5X)
2D
classification
693,550 particles
Ab-initio reconstruction
Heterogenous refinement
(3X)
86%
14%
NU refinement
3.19 Å
464,408 particles
D
3.0
3.5
4.0
4.5
5.0
E
GSFSC Resolution: 3.19Å
No Mask (3.7Å)
Loose (3.3Å)
Tight (3.2Å)
Corrected (3.2Å)
1.0
0.8
0.6
0.4
0.2
0.0
DC
19Å
9.6Å
6.4Å
4.8Å
3.8Å
3.2Å
2.7Å
2.4Å
F
Elevation
Azimuth
### of images
π/2
π/4
0
−π/4
−π/2
−π
−3π/4
−π/2
−π/4
0
π/4
π/2
3π/4
π
10³
10²
10¹
10⁰

**Figure S3: Cryo-EM data processing pipeline of Apo-C3aR-Go complex (200 kV Glacios Data).**

**A.** Representative cryo-EM micrographs of Apo-C3aR-Go complex. **B.** Representative 2D class averages with distinct secondary structures and different orientations. **C.** Schematic representation of the cryo-EM data processing workflow. **D,** Local resolution map of the 3D reconstruction. **E.** Gold standard fourier shell correlation curve (GSFSC) at 0.143 threshold. **F.** Angular distribution of the particles used for final refinement.

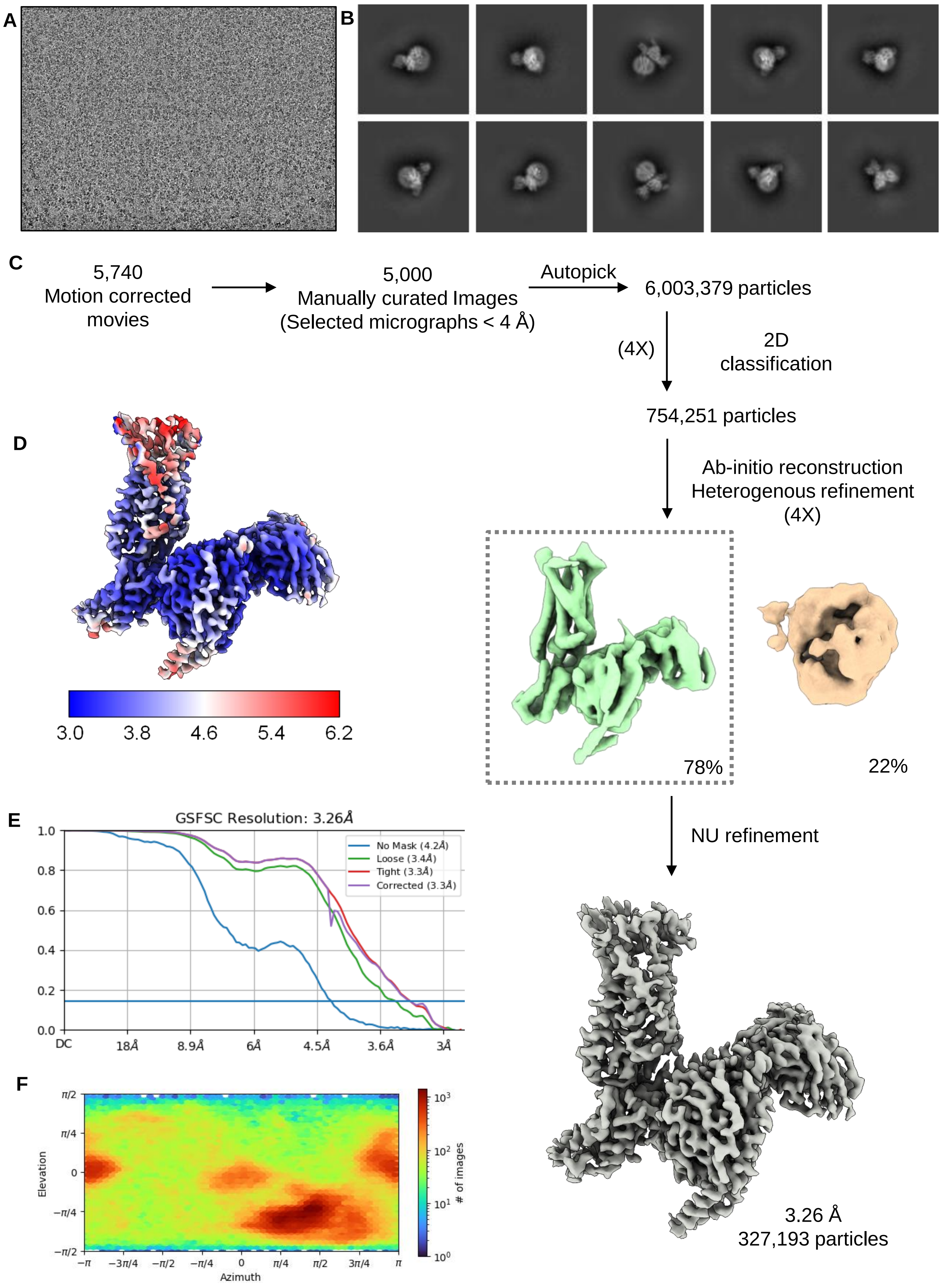
A
B
C
5,740
Motion corrected movies
5,000
Manually curated Images
(Selected micrographs < 4 Å)
Autopick
6,003,379 particles
(4X)
2D classification
754,251 particles
Ab-initio reconstruction
Heterogenous refinement
(4X)
78%
22%
NU refinement
3.26 Å
327,193 particles
D
3.0
3.8
4.6
5.4
6.2
E
GSFSC Resolution: 3.26Å
No Mask (4.2Å)
Loose (3.4Å)
Tight (3.3Å)
Corrected (3.3Å)
1.0
0.8
0.6
0.4
0.2
0.0
DC
18Å
8.9Å
6Å
4.5Å
3.6Å
3Å
F
Elevation
Azimuth
### of images
π/2
π/4
0
−π/4
−π/2
−π
−3π/4
−π/2
−π/4
0
π/4
π/2
3π/4
π
$10^3$
$10^2$
$10^1$
$10^0$

**Figure S4: Cryo-EM data processing pipeline of Apo-C3aR-Go complex (300 kV Titan Krios Data).**

**A.** Representative cryo-EM image of Apo-C3aR-Go complex. **B.** Representative 2D class averages showing distinct features and orientations of each component. **C.** Workflow of cryo-EM data processing. **D,** Cryo-EM maps are coloured by local resolution (Å). **E.** Gold standard FSC curves at 0.143 threshold. **F.** Angular distribution of the particles used for final reconstruction.

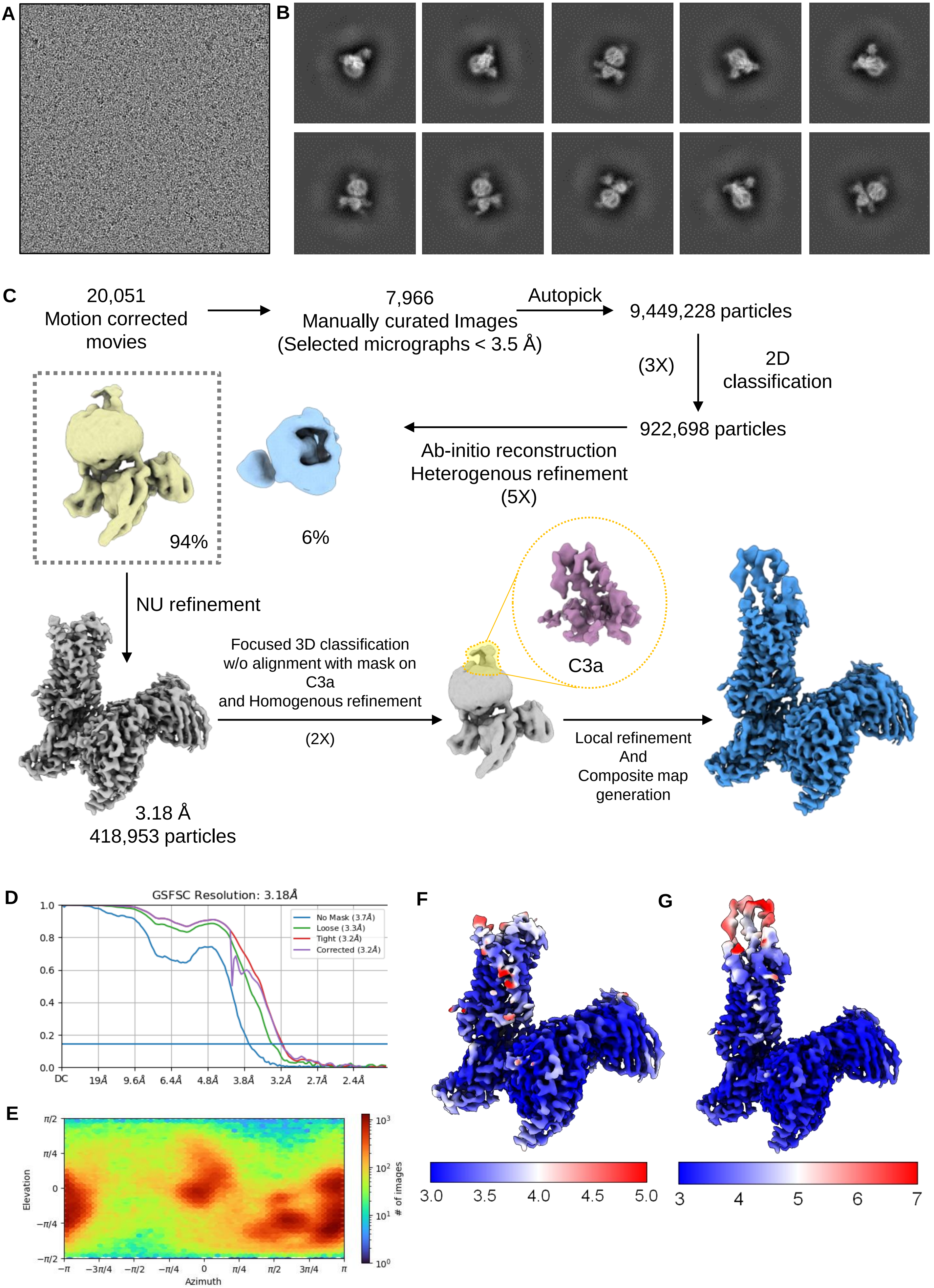
A
B
C
20,051
Motion corrected
movies
7,966
Manually curated Images
(Selected micrographs < 3.5 Å)
Autopick
9,449,228 particles
(3X)
2D
classification
922,698 particles
Ab-initio reconstruction
Heterogenous refinement
(5X)
94%
6%
NU refinement
3.18 Å
418,953 particles
Focused 3D classification
w/o alignment with mask on
C3a
and Homogenous refinement
(2X)
C3a
Local refinement
And
Composite map
generation
D
GSFSC Resolution: 3.18Å
No Mask (3.7Å)
Loose (3.3Å)
Tight (3.2Å)
Corrected (3.2Å)
1.0
0.8
0.6
0.4
0.2
0.0
DC
19Å
9.6Å
6.4Å
4.8Å
3.8Å
3.2Å
2.7Å
2.4Å
E
Elevation
Azimuth
π/2
π/4
0
−π/4
−π/2
−π
−3π/4
−π/2
−π/4
0
π/4
π/2
3π/4
π
### of images
$10^3$
$10^2$
$10^1$
$10^0$
F
3.0
3.5
4.0
4.5
5.0
G
3
4
5
6
7

**Figure S5: Cryo-EM data processing pipeline of C3a-C3aR-Go complex.**

**A.** Representative cryo-EM image of the C3a-C3aR-Go complex. **B.** Representative 2D class averages showing distinct features and orientations of each component. **C.** Cryo-EM data processing pipeline. **D,** Local resolution map of the final reconstruction. **E.** Gold standard FSC curves at 0.143 threshold. **F.** Angular distribution of the particles used for final reconstruction.

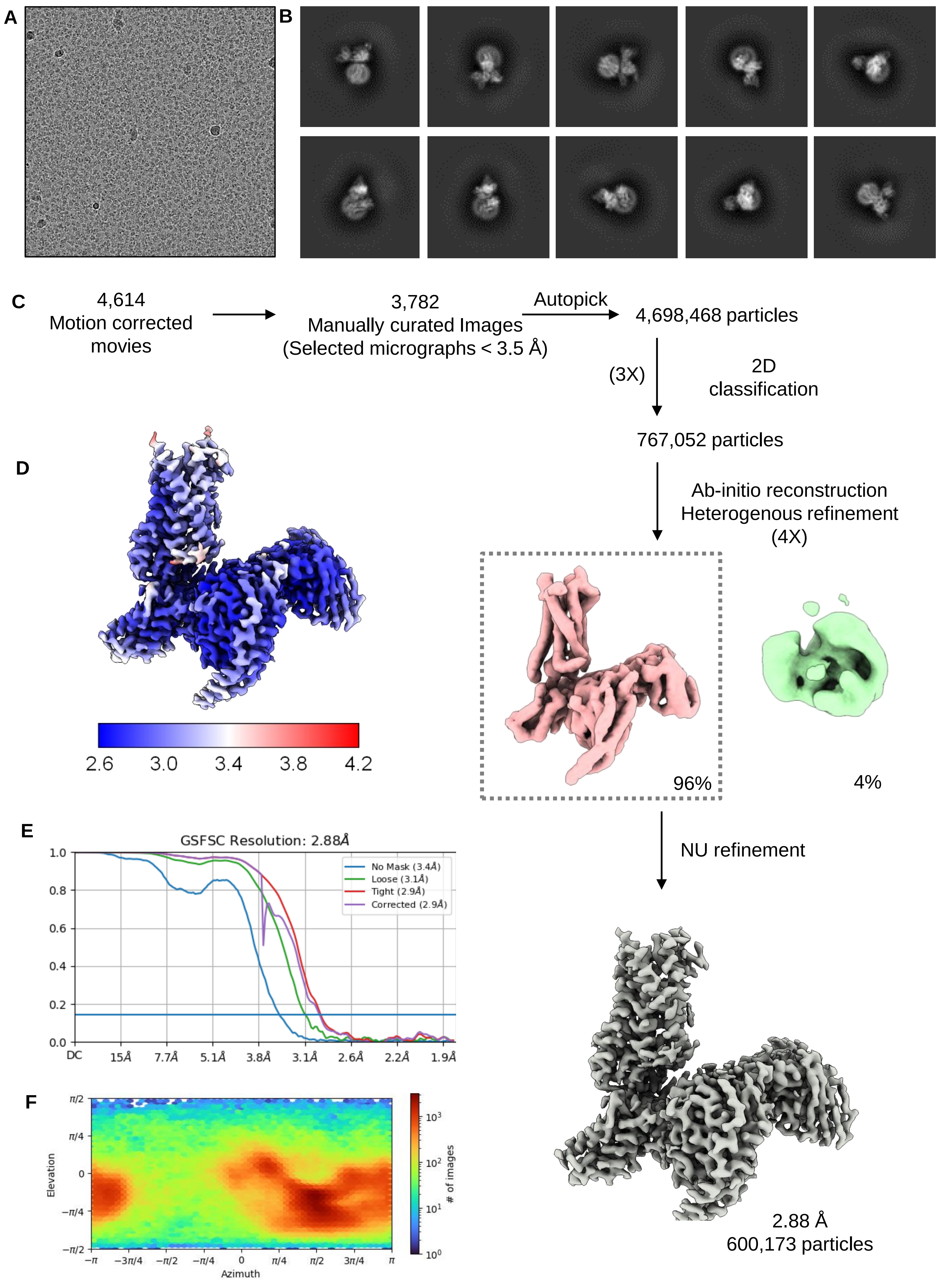

A
B
C
4,614
Motion corrected movies
3,782
Manually curated Images
(Selected micrographs < 3.5 Å)
Autopick
4,698,468 particles
(3X)
2D classification
767,052 particles
Ab-initio reconstruction
Heterogenous refinement
(4X)
96%
4%
NU refinement
2.88 Å
600,173 particles
D
2.6
3.0
3.4
3.8
4.2
E
GSFSC Resolution: 2.88Å
No Mask (3.4Å)
Loose (3.1Å)
Tight (2.9Å)
Corrected (2.9Å)
1.0
0.8
0.6
0.4
0.2
0.0
DC
15Å
7.7Å
5.1Å
3.8Å
3.1Å
2.6Å
2.2Å
1.9Å
F
Elevation
Azimuth
### of images

**Figure S6: Cryo-EM data processing pipeline of EP54-C3aR-Go complex.**

**A.** Representative cryo-EM image of the EP54-C3aR-Go complex. **B.** Representative 2D class averages showing distinct features and orientations of each component. **C.** Flowchart of the cryo-EM data processing pipeline. **D.** Local resolution map of the final reconstruction. **E.** Gold standard fourier shell correlation (GSFSC) curves at 0.143 threshold. **F.** Angular distribution of the particles used for final reconstruction.

**Data collection and processing**

| | Apo-C3aR-Go PDB-8I9S EMD-35282 | Apo-C3aR-Go PDB-8I97 EMD-35259 | C3a-C3aR-Go PDB-8I9L EMD-35275 | EP54-C3aR-Go PDB-8I95 EMD-35257 | EP54-C3aR-Gq PDB-8I9A EMD-35263 |
|---|---|---|---|---|---|
| Microscope | Titan Krios | Glacios | Glacios | Glacios | Glacios |
| Camera | GIF/K3 | Falcon 4 | Falcon 4 | Falcon 4 | Falcon 4 |
| Magnification | 105,000 | 150,000 | 150,000 | 150,000 | 150,000 |
| Voltage (kV) | 300 | 200 | 200 | 200 | 200 |
| Defocus range (µm) | -0.8 to -3.0 | -0.8 to -3.0 | -0.8 to -3.0 | -0.8 to -3.0 | -0.8 to -3.0 |
| Total dose ($e^-/A^2$) | 50 | 50 | 50 | 50 | 50 |
| Pixel size (Å) | 0.86 | 0.92 | 0.92 | 0.92 | 0.92 |
| Micrographs (no.) | 5,740 | 4,611 | 20,051 | 4,614 | 4,445 |
| Initial particles (no.) | 6,003,379 | 4,543,010 | 9,449,228 | 4,698,468 | 2,007,547 |
| Symmetry imposed | C1 | C1 | C1 | C1 | C1 |
| Final particles (no.) | 327,193 | 464,408 | 418,953 | 600,173 | 101,400 |
| Map resolution (Å) | 3.26 | 3.19 | 3.18 | 2.88 | 3.57 |
| FSC threshold | 0.143 | 0.143 | 0.143 | 0.143 | 0.143 |
| **Refinement** | | | | | |
| Initial model (PDB Code) | 8I97 | 8I95 | 8I95 | 8HPT | 8I95 |
| Model resolution (Å) | 3.5 | 3.4 | 3.5 | 3.0 | 3.8 |
| FSC threshold | 0.5 | 0.5 | 0.5 | 0.5 | 0.5 |
| **Model composition** | | | | | |
| Non-hydrogen atoms | 8,130 | 8,130 | 8,616 | 8,219 | 8,410 |
| Protein residues | 1113 | 1113 | 1183 | 1126 | 1130 |
| Ligand atoms | 0 | 0 | 0 | 1 (DAL) | 1 (DAL) |
| **R. M.S. deviations** | | | | | |
| Bond length (Å) | 0.01 | 0.005 | 0.04 | 0.004 | 0.008 |
| Bond angle (º) | 0.919 | 1.028 | 0.994 | 0.98 | 0.881 |
| **Validation** | | | | | |
| Favored (%) | 95.34 | 94.79 | 95.44 | 94.95 | 96.46 |
| Allowed (%) | 4.66 | 5.21 | 4.56 | 5.05 | 3.54 |
| Disallowed (%) | 0 | 0 | 0 | 0 | 0 |
| MolProbity score | 1.71 | 1.79 | 1.79 | 1.64 | 1.68 |
| Clash Score | 7 | 7.82 | 8.8 | 5.40 | 7.05 |

**Figure S7: Cryo-EM data collection, processing, model refinement and validation statistics.**

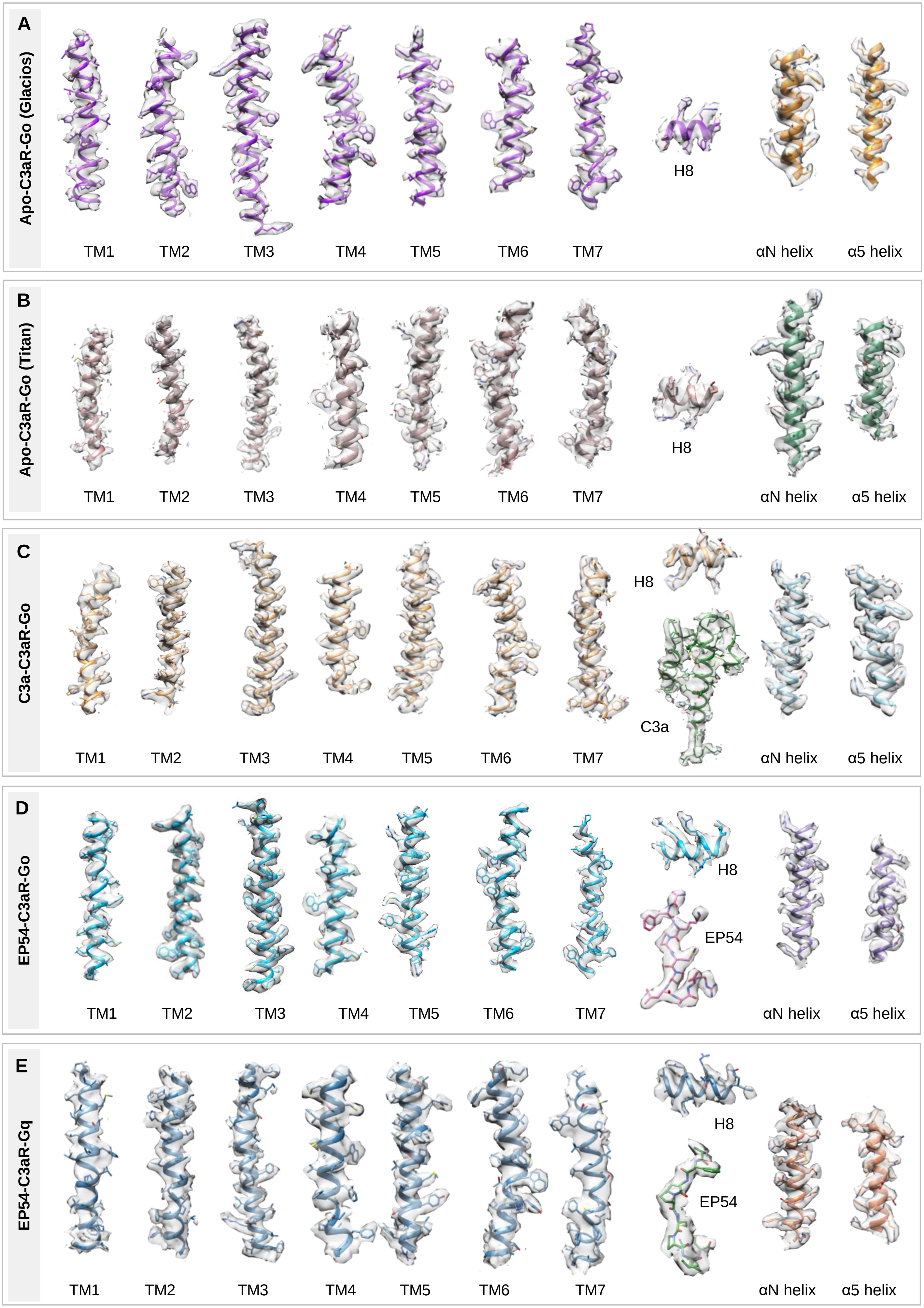
A
Apo-C3aR-Go (Glacios)
TM1
TM2
TM3
TM4
TM5
TM6
TM7
H8
αN helix
α5 helix
B
Apo-C3aR-Go (Titan)
TM1
TM2
TM3
TM4
TM5
TM6
TM7
H8
αN helix
α5 helix
C
C3a-C3aR-Go
TM1
TM2
TM3
TM4
TM5
TM6
TM7
H8
C3a
αN helix
α5 helix
D
EP54-C3aR-Go
TM1
TM2
TM3
TM4
TM5
TM6
TM7
H8
EP54
αN helix
α5 helix
E
EP54-C3aR-Gq
TM1
TM2
TM3
TM4
TM5
TM6
TM7
H8
EP54
αN helix
α5 helix

#### Figure S8: Representative cryo-EM maps.

**A-E.** Cryo-EM densities corresponding to the TMs and helix 8 of the receptor, αN and α5 of Go, Gq and Ligands are shown. Complex names have been mentioned in the left inside respective boxes.

| Component | Total residues | Resolved residues | Total residues | Resolved residues | Total residues | Resolved residues | Total residues | Resolved residues | Total residues | Resolved residues |
|---|---|---|---|---|---|---|---|---|---|---|
| | **C3a-C3aR-Go** | | **EP54-C3aR-Go** | | **EP54-C3aR-Gq** | | **Apo-C3aR-Go (Glacios)** | | **Apo-C3aR-Go (Titan)** | |
| **C3a/ EP54** | S1-R77 | S1-G26<br>F34-R77 | A1-R10 | S2-R10 | A1-R10 | S2-R10 | NA | NA | NA | NA |
| **C3aR** | M1-V482 | W18$_{N\text{-}term}$-Y174$_{ECL2}$<br>T331$_{5.3}$ I450$^{C\text{-}}_{term}$ | M1-V482 | P17$_{N\text{-}term}$-Y174$_{ECL2}$<br>T331$_{5.33}$-G452$_{C\text{-}term}$ | M1-V482 | P17$_{N\text{-}term}$-Y174$_{ECL2}$<br>T331$_{5.33}$-M363$_{ICL3}$<br>Q372-G452$_{C\text{-}term}$ | M1-V482 | N19$_{N\text{-}term}$-Y174$_{ECL2}$<br>T331$_{5.33}$-I450$_{C\text{-}term}$ | M1-V482 | N19$_{N\text{-}term}$-Y174$_{ECL2}$<br>T331$_{5.33}$-I450$_{C\text{-}term}$ |
| **mini Gαo/ Gαq** | M1-H57<br>T172 - Y366 | S6-I55<br>T183-Y231<br>R243-Y354 | M1-H57<br>T172-Y366 | S6-I55<br>T183-Y354 | M1-V359 | E15-I62<br>T204-D252<br>N264-T364<br>D368-V394 | M1-H57<br>T172-Y366 | S6-I55<br>T183-Y231<br>R243-Y354 | M1-H57<br>T172-Y366 | S6-I55<br>T183-D230<br>R243-Y354 |
| **Gβ** | M1-N340 | L4-N340 | M1-N340 | E3-N340 | M1-N340 | E3-N340 | M1-N340 | L4-N340 | M1-N340 | E3-N340 |
| **Gγ** | M1-L71 | S8-R62 | M1-L71 | A7-R62 | M1-L71 | A7-R62 | M1-L71 | S8-R62 | M1-L71 | A7-R62 |
| **ScFv16** | D1-K248 | D1-S120<br>G135-K248 | D1-K248 | D1-S120<br>G135-K248 | D1-K248 | D1-R72<br>K76-S120<br>G135-K248 | D1-K248 | D1-S120<br>G135-K248 | D1-K248 | D1-S120<br>G135-K248 |

**Figure S9: List of residues resolved in the structures of C3a-C3aR-Go, EP54-C3aR-Go, EP54-C3aR-Gq and Apo-C3aR-Go.**

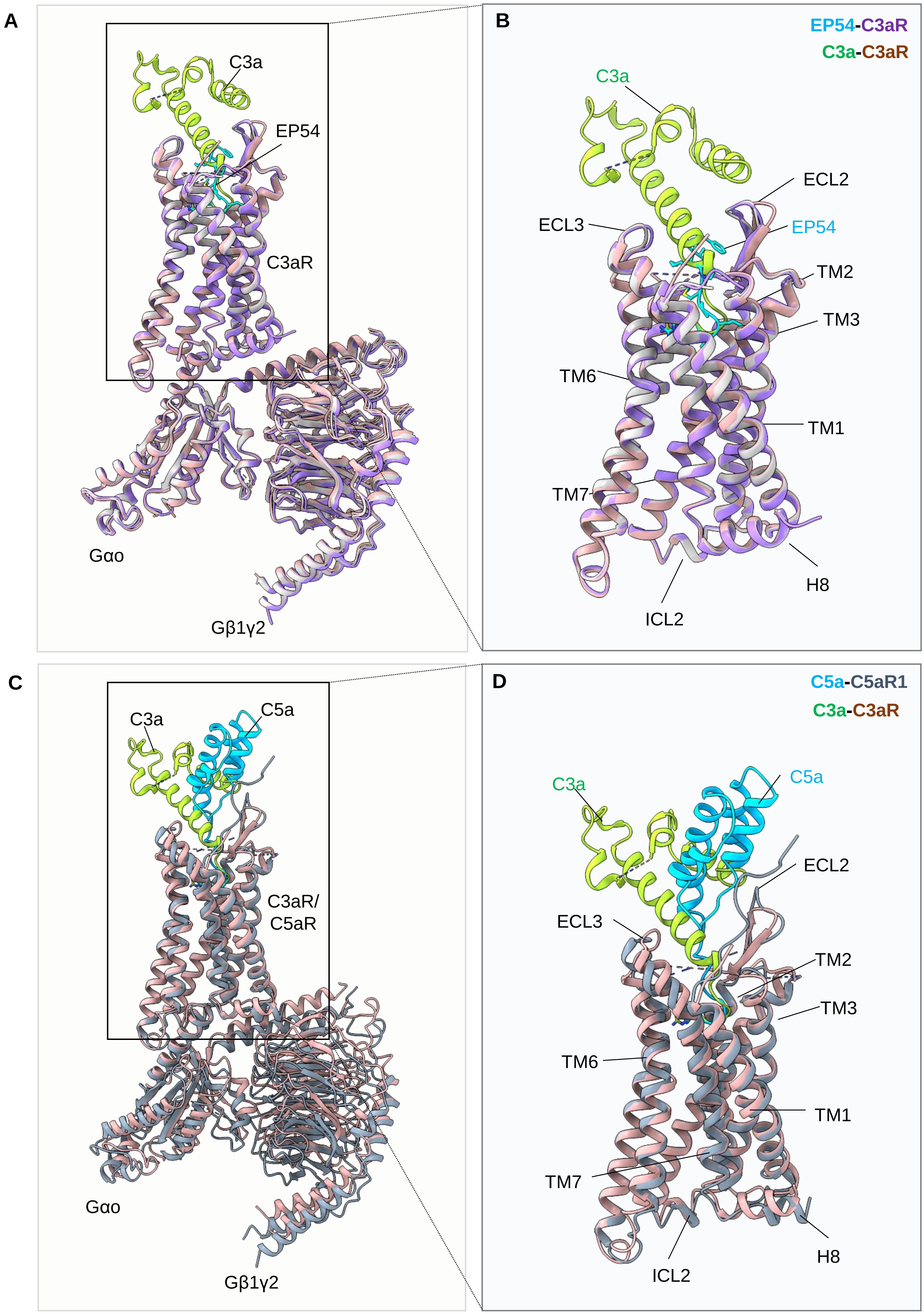
A
C3a
EP54
C3aR
Gαo
Gβ1γ2
B
EP54-C3aR
C3a-C3aR
C3a
ECL2
ECL3
EP54
TM2
TM3
TM6
TM1
TM7
H8
ICL2
C
C3a
C5a
C3aR/
C5aR
Gαo
Gβ1γ2
D
C5a-C5aR1
C3a-C3aR
C3a
C5a
ECL2
ECL3
TM2
TM3
TM6
TM1
TM7
H8
ICL2

**Figure S10: Structural alignment of C3aR and C5aR1 structures.**

**A.** Alignment of the EP54/C3a-C3aR-Go complexes. **B.** Alignment of ligand bound C3aR without Go. **C.** Alignment of the C3a-C3aR-Go and C5a-C5aR1-Go complexes. **D.** Alignment of ligand bound C3aR and C5aR1 without Go.

A

| C3a-C3aR interface | | |
|---|---|---|
| **Chain C (C3aR)** | **Distance (Å)** | **Chain D (C3a)** |
| Phe77 (TM2) | Leu75 (3.62) | Leu75 |
| Ser78 (TM2) | Leu75 (3.62) | Leu75 |
| Trp88 (ECL1) | Leu75 (3.74) | Leu75 |
| Pro99 (TM3) | Leu75 (3.63) | Leu75 |
| Ile102 (TM3) | Leu75 (3.46) | Leu75 |
| Val157 (TM4) | Arg77 (3.87) | Arg77 |
| Glu162 (ECL2) | Gln66 (2.96) | Gln66 |
| Phe164 (ECL2) | Gln66 (3.09), His67 (3.81) | Gln66, His67 |
| Thr166 (ECL2) | Gln3 (3.12) | Gln3 |
| Asp167 (ECL2) | Gln3 (3.84) | Gln3 |
| Arg171 (ECL2) | Ala70 (3.41), Ser71 (3.45) | Ala70, Ser71 |
| Tyr174 (ECL2) | Arg77 (2.33) | Arg77 |
| Arg340 (TM5) | Ala76 (3.74), Arg77 (3.55) | Ala76, Arg77 |
| Tyr393 (TM6) | Ala76 (3.73), Arg77 (2.94) | Ala76, Arg77 |
| Phe396 (TM6) | Arg77 (3.77) | Arg77 |
| Gly397 (TM6) | Arg77 (3.11) | Arg77 |
| Pro405 (ECL3) | Arg65 (3.57) | Arg65 |
| Asp417 (TM7) | Arg77 (2.64) | Arg77 |
| His418 (TM7) | Gly74 (3.80) | Gly74 |
| Ile421 (TM7) | Ala76 (3.51) | Ala76 |

B

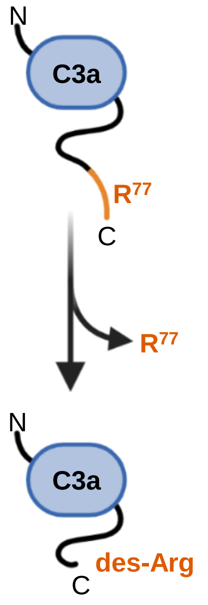


C

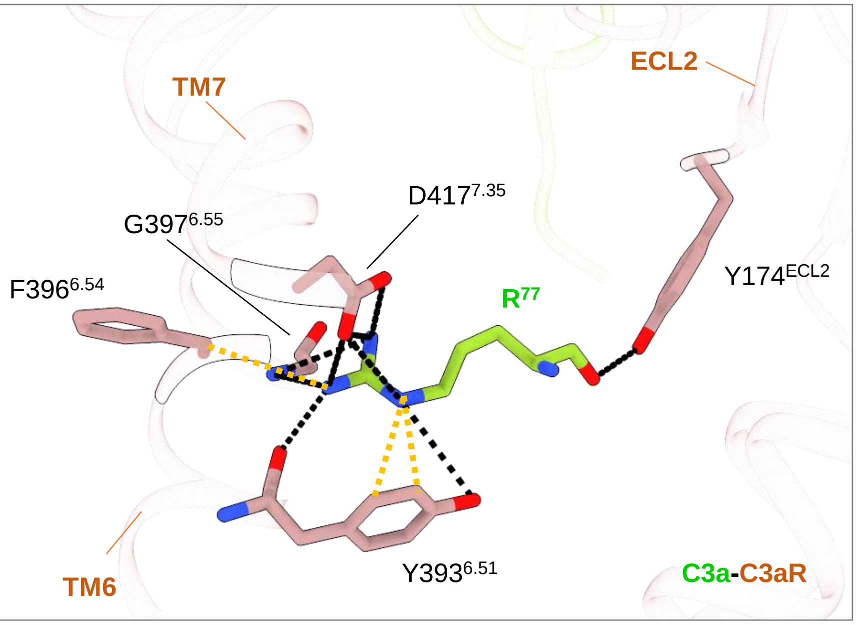

**Figure S11: Interactions of C3a with C3aR and C3a-desArg.**

**A.** Residues within 4 Å radius are shown. **B.** Proteolytic cleavage of the C-terminal $Arg^{77}$ from C3a results in C3a-desArg. **C.** Interactions of $Arg^{77}$ with the residues of C3aR in the orthosteric pocket are shown.

**A**

| EP54-C3aR$^{Go}$ interface | | |
|---|---|---|
| **Chain C (C3aR)** | **Distance (Å)** | **Chain D (EP54)** |
| Phe77 (TM2) | Leu8 (3.65) | Leu8 |
| Ser78 (TM2) | Leu8 (3.64) | Leu8 |
| His81 (TM2) | Leu8 (3.79) | Leu8 |
| Trp88 | Leu8 (3.69) | Leu8 |
| Pro99 | Leu8 (3.21) | Leu8 |
| Ile102 (TM3) | Leu8 (3.61), Dal9 (3.31) | Leu8, Dal9 |
| Val103 (TM3) | Dal9 (3.36), Arg10 (3.79) | Dal9, Arg10 |
| Met106 (TM3) | Dal9 (3.79) | Dal9 |
| Val157 | Arg10 (3.78) | Arg10 |
| Arg161 (TM4) | Pro7 (3.84), Arg10 (3.75) | Pro7, Arg10 |
| Glu162 (ECL2) | Phe3 (3.55) | Phe3 |
| Phe164 (ECL2) | Phe3 (3.38) | Phe3 |
| Arg171 (ECL2) | Phe3 (3.01) | Phe3 |
| Cys172 (ECL2) | Phe3 (3.72) | Phe3 |
| Gly173 (ECL2) | Phe3 (3.19) | Phe3 |
| Tyr174 (ECL2) | Ser2 (2.95), Phe3 (3.87), Lys4 (3.15), Met6 (3.18), Arg10 (2.68) | Ser2, Phe3, Lys4, Met6, Arg10 |
| Leu333 (TM5) | Lys4 (3.54), Met6 (3.37) | Lys4, Met6 |
| Arg340 (TM5) | Arg10 (2.40) | Arg10 |
| Tyr393 (TM6) | Arg10 (2.95), Dal9 (2.94) | Arg10, Dal9 |
| Phe396 (TM6) | Arg10 (3.39) | Arg10 |
| Gly397 (TM6) | Arg10 (3.16) | Arg10 |
| Asp417 (TM7) | Pro7 (3.31), Arg10 (2.70) | Pro7, Arg10 |
| Cys420 (TM7) | Arg10 (3.83) | Arg10 |
| Ile421 (TM7) | Pro7 (3.58), Arg10 (3.83) | Pro7, Arg10 |

**Figure S12: Interactions of EP54 with C3aR.** Residues within 4 Å radius are shown.

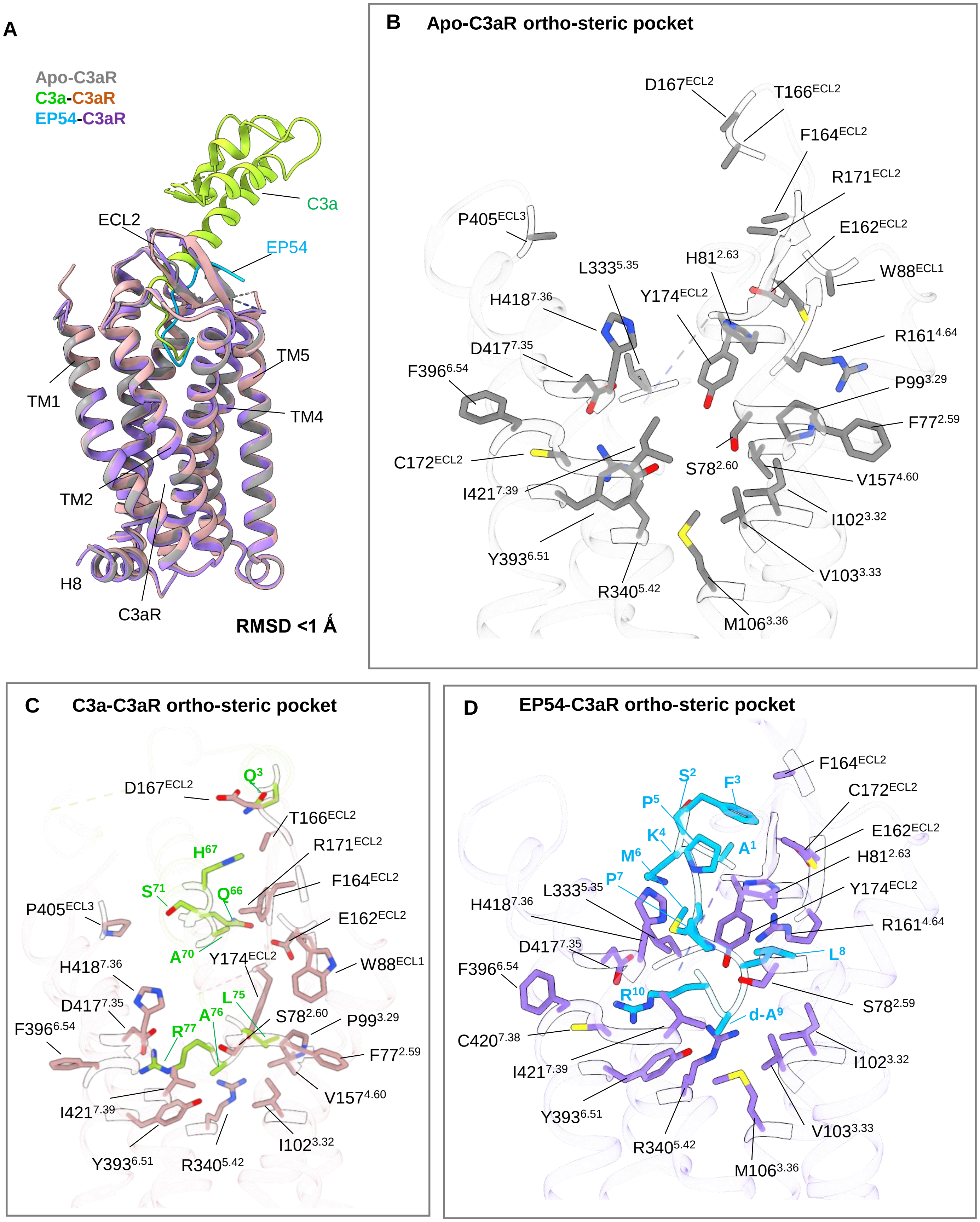

A
Apo-C3aR
C3a-C3aR
EP54-C3aR
C3a
ECL2
EP54
TM5
TM1
TM4
TM2
H8
C3aR
RMSD <1 Å
B
Apo-C3aR ortho-steric pocket
D167$^{ECL2}$
T166$^{ECL2}$
F164$^{ECL2}$
R171$^{ECL2}$
P405$^{ECL3}$
E162$^{ECL2}$
L333$^{5.35}$
H81$^{2.63}$
W88$^{ECL1}$
H418$^{7.36}$
Y174$^{ECL2}$
R161$^{4.64}$
D417$^{7.35}$
F396$^{6.54}$
P99$^{3.29}$
F77$^{2.59}$
C172$^{ECL2}$
S78$^{2.60}$
V157$^{4.60}$
I421$^{7.39}$
I102$^{3.32}$
Y393$^{6.51}$
V103$^{3.33}$
R340$^{5.42}$
M106$^{3.36}$
C
C3a-C3aR ortho-steric pocket
Q$^{3}$
D167$^{ECL2}$
T166$^{ECL2}$
R171$^{ECL2}$
H$^{67}$
F164$^{ECL2}$
S$^{71}$
Q$^{66}$
P405$^{ECL3}$
E162$^{ECL2}$
A$^{70}$
Y174$^{ECL2}$
W88$^{ECL1}$
H418$^{7.36}$
L$^{75}$
D417$^{7.35}$
A$^{76}$
S78$^{2.60}$
P99$^{3.29}$
F396$^{6.54}$
R$^{77}$
F77$^{2.59}$
V157$^{4.60}$
I421$^{7.39}$
I102$^{3.32}$
Y393$^{6.51}$
R340$^{5.42}$
D
EP54-C3aR ortho-steric pocket
F164$^{ECL2}$
S$^{2}$
F$^{3}$
C172$^{ECL2}$
P$^{5}$
E162$^{ECL2}$
K$^{4}$
A$^{1}$
M$^{6}$
H81$^{2.63}$
P$^{7}$
L333$^{5.35}$
Y174$^{ECL2}$
H418$^{7.36}$
R161$^{4.64}$
D417$^{7.35}$
L$^{8}$
F396$^{6.54}$
S78$^{2.59}$
R$^{10}$
d-A$^{9}$
C420$^{7.38}$
I102$^{3.32}$
I421$^{7.39}$
Y393$^{6.51}$
V103$^{3.33}$
R340$^{5.42}$
M106$^{3.36}$

**Figure S13: Comparison of the residues in the orthosteric pocket of C3aR in Apo, EP54 and C3a bound states.**

**A.** Structural superimposition of Apo, EP54 and C3a bound C3aR-Go complexes. **B.** Conformation of all the residues present in the orthosteric pocket of C3aR involved in interaction with the ligands are shown in the Apo structure. **C, D.** Residues of the orthosteric pocket of C3aR involved in interaction with the residues of C3a (**C**) and EP54 (**D**) are highlighted.

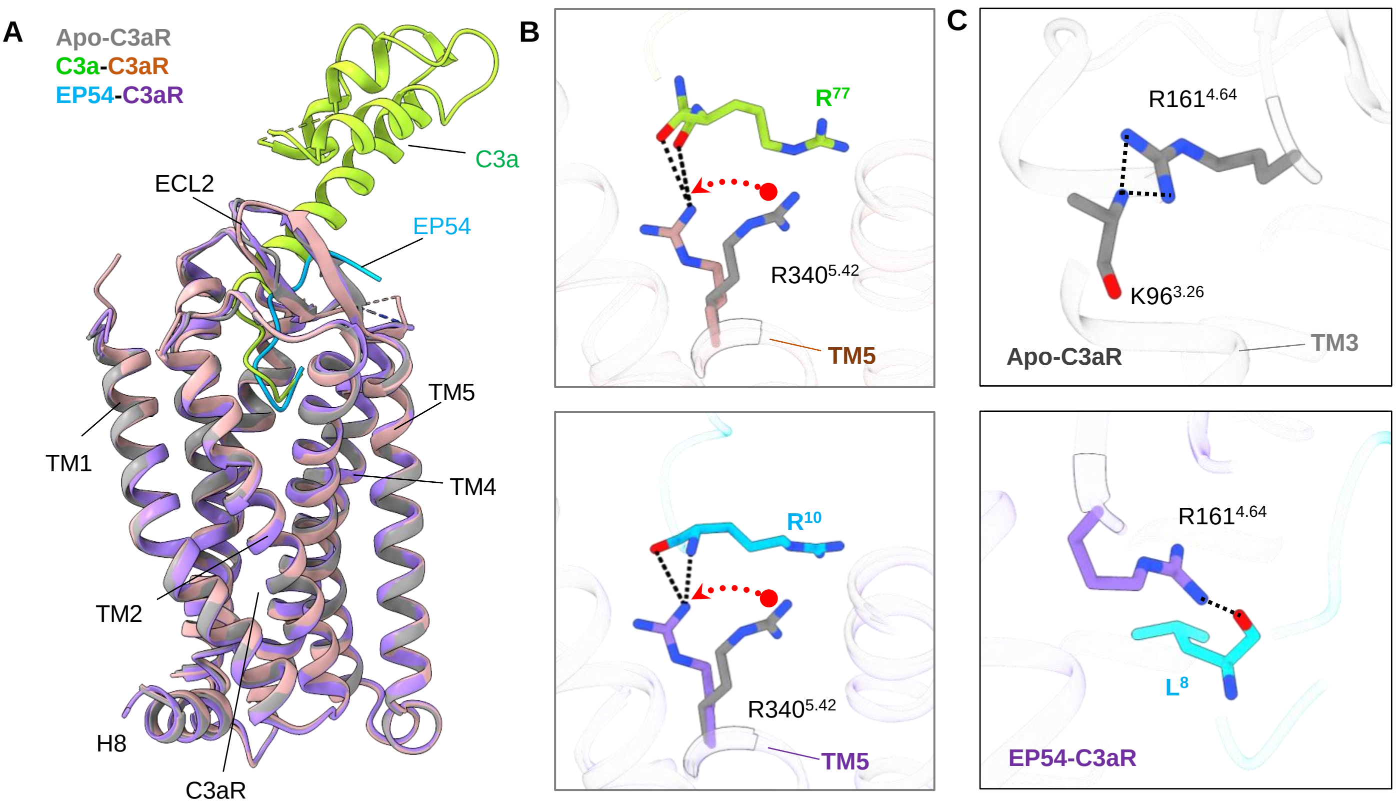
A
Apo-C3aR
C3a-C3aR
EP54-C3aR
C3a
ECL2
EP54
TM5
TM1
TM4
TM2
H8
C3aR
B
R77
R340 5.42
TM5
R10
R340 5.42
TM5
C
R161 4.64
K96 3.26
Apo-C3aR
TM3
R161 4.64
L8
EP54-C3aR

#### Figure S14: Structure of C3aR in Apo state.

**A.** Structural alignment of Apo with EP54/C3a bound C3aR. **B, C.** Changes in rotameric conformations of Arg340$^{5.42}$ and Arg161$^{4.64}$ in the Apo state as compared to the ligand bound state.

A

| C3aR$^{Apo}$-Go interface | | |
|---|---|---|
| **Chain C (C3aR)** | **Distance (Å)** | **Chain A (Go)** |
| Asn57 (TM2) | Gly350 (2.97) | Gly350 |
| Asp119 (TM3) | Cys351 (3.84) | Cys351 |
| Arg120 (TM3) | Cys351 (3.86) | Cys351 |
| Val123 (TM3) | Asn347(3.08) | Asn347 |
| Val124 (TM3) | Ile344 (3.61), Leu348 (3.62) | Ile344, Leu348 |
| Pro127 (ICL2) | Ile343 (3.58), Ile344 (3.44) | Ile343, Ile344 |
| Ile128 (ICL2) | Lys193 (3.78), Phe336 (3.43) | Lys193,Phe336 |
| Gln131 (ICL2) | Lys32 (3.65), Asp33 (3.74), Val34 (3.15), Leu195 (3.59), Ile343 (3.19), | Lys32, Val34, Leu195, Ile343, Asp33 |
| Asn132 (ICL2) | Lys32 (2.41), Asn194 (3.01) | Lys32, Asn194 |
| Asn135 (TM4) | Ile28(3.43) | Ile28 |
| Ile359 (TM5) | Leu353(3.61) | Leu353 |
| Arg362 (TM5) | Ile344(3.27) | Ile344 |
| Arg367 (ICL3) | Ser264 (3.88), Asn316 (3.46) Glu318 (3.18), Tyr320 (3.32), Tyr354 (3.21), Ala345 (3.25), Arg349 (3.16) | Tyr320, Tyr354, Glu318, Ala345, Ser264, Asn316, Arg349 |
| Phe368 (ICL3) | Ile344 (3.47), Ala345 (3.78), Tyr354 (3.28) | Ile344, Ala345, Tyr354 |
| Ser371 (ICL3) | Tyr354 (3.22) | Tyr354 |
| Lys374 (TM6) | Tyr354 (3.60) | Tyr354 |
| Thr375 (TM6) | Leu353 (2.55), Tyr354 (3.88) | Leu353, Tyr354 |
| Val378 (TM6) | Leu353 (3.71) | Leu353 |

**Figure S15: Interactions of Go with $C3aR^{Apo}$.** Residues within 4 Å radius are shown.

A

| C3aR$^{C3a}$-Go interface | | |
|---|---|---|
| **Chain C (C3aR)** | **Distance (Å)** | **Chain A (Go)** |
| Asn135 (TM4) | Ile28(3.41) | Ile28 |
| Gln131 (ICL2) | Lys32 (3.80), Val34 (3.25), Leu195 (3.22), Ile343 (3.20) | Lys32 , Val34, Leu195, Ile343 |
| Asn132 (ICL2) | Lys32 (3.24), Asn194 (2.73) | Lys32, Asn194 |
| Arg367 (ICL3) | Asn316 (3.77), Glu318 (3.21), Tyr320 (2.91), Asp341 (3.46), Ala345 (3.64) , Tyr354 (3.05) | Asn136, Glu318, Tyr320, Asp341, Ala345, Tyr354 |
| Pro127 (ICL2) | Ile343 (3.89), Ile344 (3.61) | Ile343, Ile344 |
| Val124 (TM3) | Ile344 (3.28), Leu348 (3.65) | Ile344, Leu348 |
| Arg362 (TM5) | Ile344 (3.40) | Ile344 |
| Phe368 (ICL3) | Ala345 (3.89), Asp341(3.22), Tyr354 (3.31) | Ala345 |
| Val123 (TM3) | Asn347 (3.34) | Asn347 |
| Asn57 (TM2) | Gly350 (2.73) | Gly350 |
| Arg120 (TM3) | Cys351 (3.30) | Cys351 |
| Asp119 (TM3) | Cys351 (3.81) | Cys351 |
| Leu438 (TM7) | Gly352 (3.34) | Gly352 |
| Lys374 (TM6) | Gly352 (3.73) | Gly352 |
| Thr375 (TM6) | Leu353 (2.32),Tyr354 (3.84) | Leu353,Tyr354 |
| Tyr356 (TM5) | Leu353 (3.82) | Leu353 |
| Ile359 (TM5) | Leu353 (3.62) | Leu353 |
| Ser371 (ICL3) | Tyr354 (3.08) | Tyr354 |

**Figure S16: Interactions of Go with C3aR$^{C3a}$.** Residues within 4 Å radius are shown.

**A**

| C3aR$^{EP54}$-Go interface | | |
|---|---|---|
| **Chain C (C3aR)** | **Distance (Å)** | **Chain A (Go)** |
| Asn57 (TM2) | Gly350 (3.66) | Gly350 |
| Arg120 (TM3) | Cys351 (3.07), Leu353 (3.87) | Cys351, Leu353 |
| Val123 (TM3) | Ile344 (3.83), Asn347 (3.66) | Ile344, Asn347 |
| Val124 (TM3) | Leu348 (3.62) | Leu348 |
| Pro127 (ICL2) | Ile343 (3.72), Ile344 (3.44) | Ile343, Ile344 |
| Asn135 (TM4) | Ile28 (3.41) | Ile28 |
| Gln131 (ICL2) | Lys32 (3.82), Val34 (3.20), Leu195 (3.28), Ile343 (3.23) | Lys32 Val34, Leu195, Ile343 |
| Asn132 (ICL2) | Lys32 (2.78), Asn194 (2.85) | Lys32, Asn194 |
| Ile359 (TM5) | Leu353 (3.61) | Leu353 |
| Arg362 (TM5) | Ile344 (3.22) | Ile344 |
| Arg367 (ICL3) | Glu318 (3.11), Tyr320 (2.49), Asp341 (3.25), Ile342 (3.35), Ala345 (3.70), Arg349 (3.44), Tyr354 (2.96) | Glu318, Tyr320, Asp341, Ile342, Ala345, Ile342, Tyr354 |
| Phe368 (ICL3) | Asp341 (3.71), Ile344 (3.50), Tyr354 (3.39) | Asp341, Ile344, Tyr354 |
| Ser371 (ICL3) | Tyr354 (3.09) | Tyr354 |
| Lys374 (TM6) | Gly352 (3.06) | Gly352 |
| Thr375 (TM6) | Leu353 (2.35) | Leu353 |
| Val378 | Leu353 (3.68) | Leu353 |

**Figure S17: Interactions of Go with C3aR$^{EP54}$.** Residues within 4 Å radius are shown.

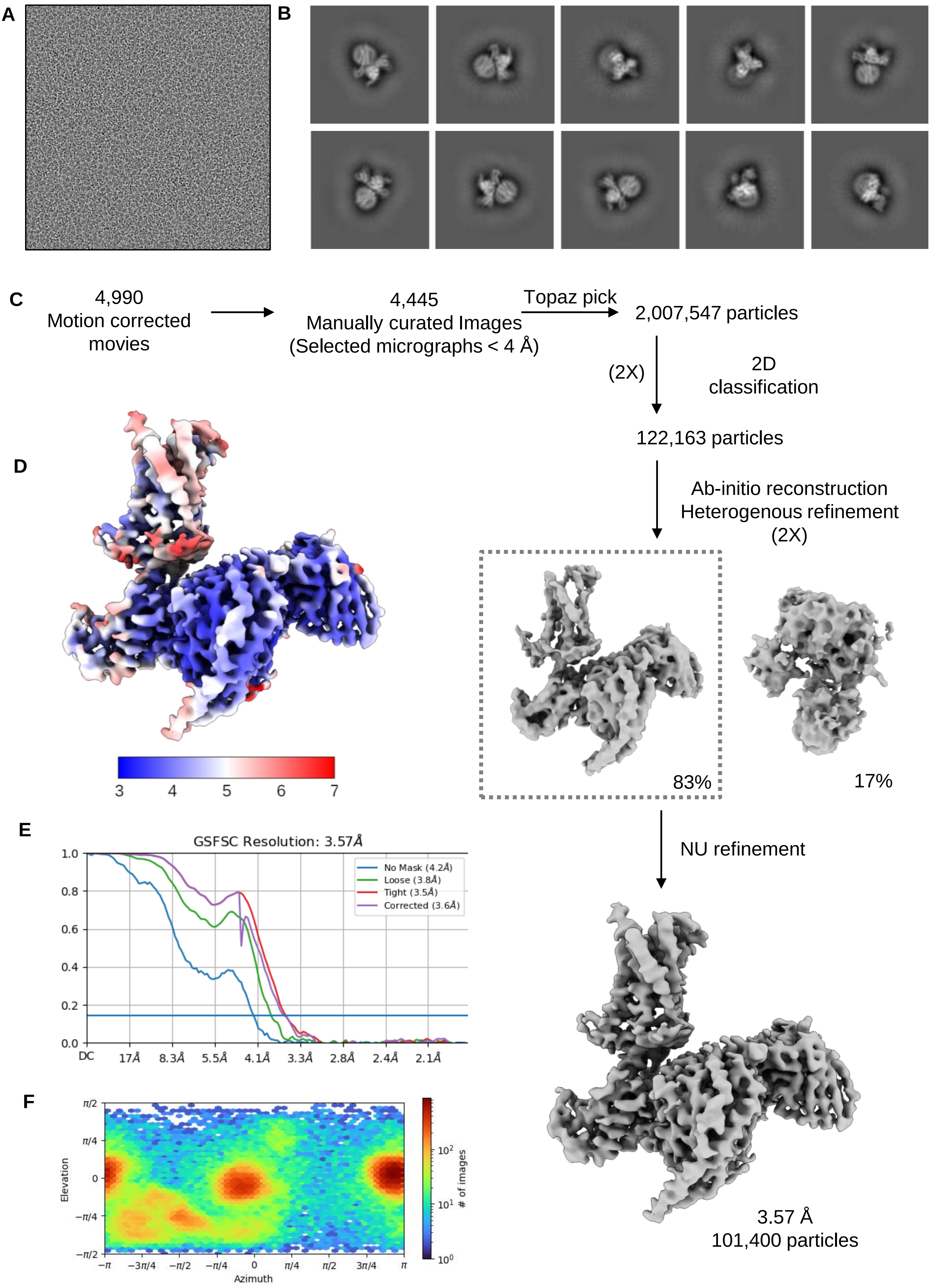

A
B
C
4,990
Motion corrected movies
4,445
Manually curated Images
(Selected micrographs < 4 Å)
Topaz pick
2,007,547 particles
(2X)
2D classification
122,163 particles
Ab-initio reconstruction
Heterogenous refinement
(2X)
83%
17%
NU refinement
3.57 Å
101,400 particles
D
3
4
5
6
7
E
GSFSC Resolution: 3.57Å
No Mask (4.2Å)
Loose (3.8Å)
Tight (3.5Å)
Corrected (3.6Å)
DC
17Å
8.3Å
5.5Å
4.1Å
3.3Å
2.8Å
2.4Å
2.1Å
F
Elevation
Azimuth
### of images

**Figure S18: Cryo-EM data processing pipeline of EP54-C3aR-Gq complex.**

**A.** Representative cryo-EM image of the EP54-C3aR-Gq complex. **B.** Representative 2D class averages showing distinct features and orientations of each component. **C.** Flowchart of the cryo-EM data processing pipeline. **D.** Local resolution map of the final reconstruction. **E.** Gold standard fourier shell correlation (GSFSC) curves at 0.143 threshold. **F.** Angular distribution of the particles used for final reconstruction.

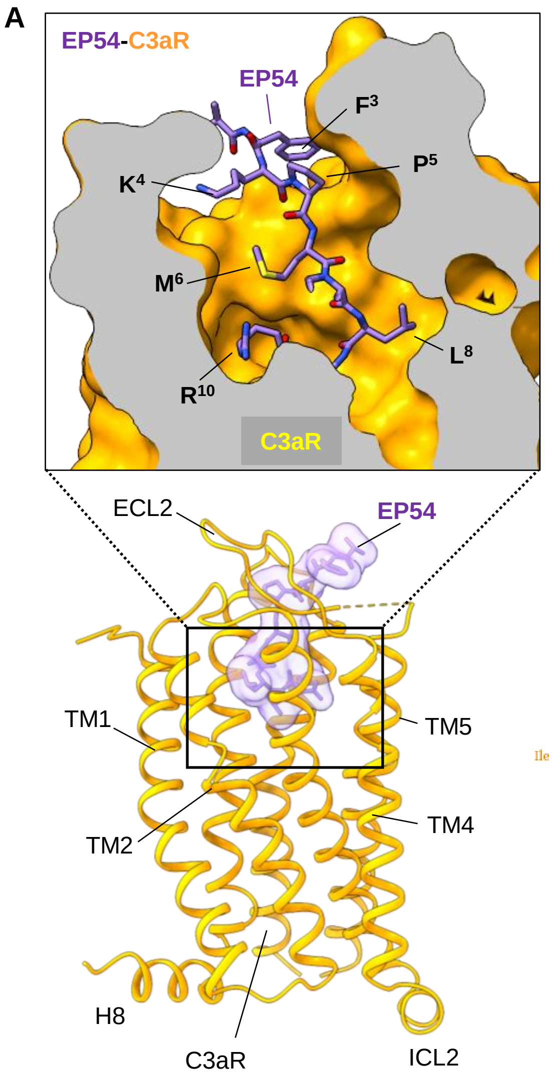


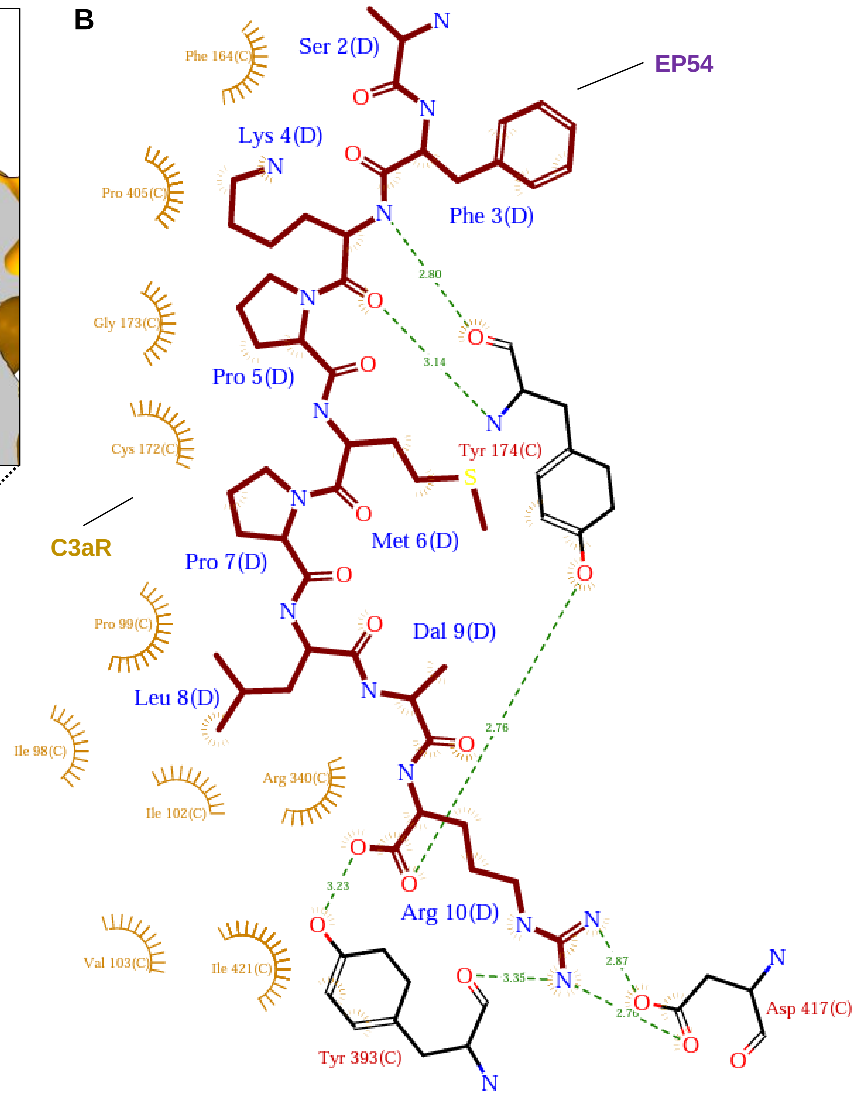


**D**

| EP54-C3aR-Go | | EP54-C3aR-Gq |
|---|---|---|
| Specific contacts | Common contacts | Specific contacts |
| F77 | P99 | I98 |
| S78 | I102 | P405 |
| H81 | V103 | |
| M106 | R161 | |
| V157 | F164 | |
| R171 | C172 | |
| L333 | G173 | |
| F396 | Y174 | |
| G397 | R340 | |
| C420 | Y393 | |
| | D417 | |
| | I421 | |

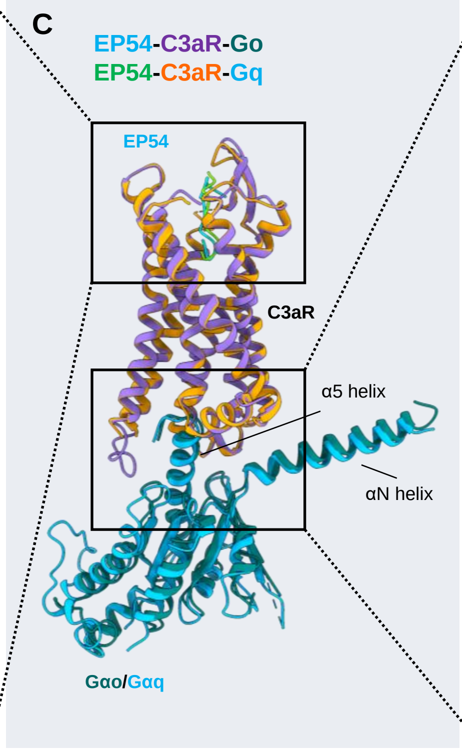


**E**

| EP54-C3aR-Go | | EP54-C3aR-Gq |
|---|---|---|
| Specific contacts | Common contacts | Specific contacts |
| D119 | N57 | R362 |
| Y435 | R120 | R367 |
| G439 | V123 | F368 |
| | V124 | S371 |
| | P127 | V378 |
| | Q131 | |
| | N132 | |
| | N135 | |
| | I359 | |
| | K374 | |
| | T375 | |

**Figure S19: EP54 binding and activation of C3aR in complex with Gq and comparison with Go bound state.**

**A.** EP54 binding pose at the orthosteric pocket of C3aR in complex Gq. EP54 in transparent surface (bottom) and receptor in surface slice showing side chains of EP54 in the ligand binding pocket (top). **B.** Interaction interface between EP54 and C3aR-Gq. The interaction plot has been generated through PDBSum. **C.** Structural superposition of EP54-C5aR-Gαo and EP54-C3aR-Gαq complexes. **D, E**. Common and specific residue interactions between C3aR and EP54 **(D)** and, C3aR and Gαo/Gαq **(E)** respectively.

**A**

| EP54-C3aR$^{Gq}$ | | |
|---|---|---|
| **Chain C (C3aR)** | **Distance (Å)** | **Chain D (EP54)** |
| Ile98 | Leu8 (3.78) | Leu8 |
| Pro99 | Leu8 (3.68) | Leu8 |
| Ile102 | Leu8 (3.67) | Leu8 |
| Val103 | Dal9 (3.59) | Dal9 |
| Arg161 | Pro7 (3.81), Leu8 (3.44), Arg10 (3.78) | Pro7, Leu8, Arg10 |
| Phe164 | Phe3 (3.11) | Phe3 |
| Cys172 | Pro5 (3.54), Met6 (3.50) | Pro5, Met6 |
| Gly173 | Lys4 (3.57) | Lys4 |
| Tyr174 | Phe3 (3.39), Lys4 (2.80), Met6 (3.63), Arg10 (2.76) | Phe3, Lys4, Met6, Arg10 |
| Arg340 | Arg10 (2.29) | Arg10 |
| Tyr393 | Dal9 (3.41), Arg10 (3.23) | Dal9 Arg10 |
| Pro405 | Lys4 (3.43) | Lys4 |
| Asp417 | Pro7 (3.75), Arg10 (2.76) | Pro7, Arg10 |
| Ile421 | Dal9 (3.04), Arg10 (3.42) | Dal9, Arg10 |

**B**

| C3aR$^{EP54}$-Gq | | |
|---|---|---|
| Chain C (C3aR) | Distance (Å) | Chain A (Gq) |
| Asn57 | Glu390 (3.62), Tyr 391 (3.12) | Glu390, Tyr 391 |
| Trp60 | Tyr391 (3.25) | Tyr 391 |
| Phe61 | Tyr391 (3.77) | Tyr 391 |
| Asp119 | Tyr391 (2.45) | Tyr 391 |
| Arg120 | Tyr391 (3.03), Leu393 (3.75) | Tyr391, Leu393 |
| Val123 | Asn387 (3.06) | Asn387 |
| Val124 | Leu384 (3.76), Leu388 (3.48) | Leu384, Leu388 |
| Pro127 | Lys380 (3.71) | Lys380 |
| Gln131 | Arg39 (2.81), Leu41 (3.35), Val217 (3.23), Ile383 (3.06) | Arg39, Leu41, Val217, Ile383 |
| Asn132 | Lys216 (3.09) | Lys216 |
| Asn135 | Gln35 (3.68), | Gln35 |
| Ile359 | Val394 (3.84) | Val394 |
| Lys374 | Asn392 (3.44), Val394 (2.90) | Asn392, Val394 |
| Thr375 | Leu393 (2.90), Val394 (3.47) | Leu393, Val394 |
| Tyr435 | Asn392 (3.65) | Asn392 |
| Gly439 | Asn392 (3.22) | Asn392 |

**Figure S20:** **A.** Interactions between EP54 and C3aR bound to Gq. **B.** Interface between C3aR and Gq. Residues within 4 Å radius are shown.

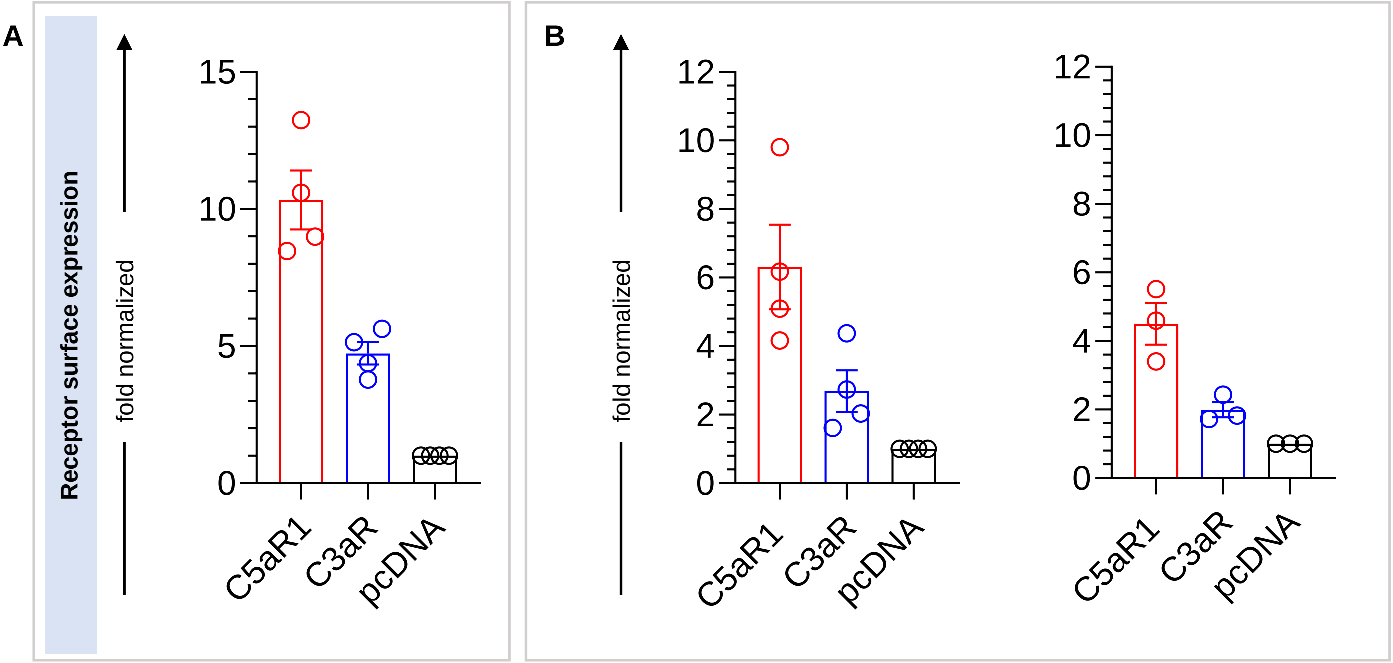
A
B
Receptor surface expression
fold normalized
fold normalized
15
10
5
0
12
10
8
6
4
2
0
12
10
8
6
4
2
0
C5aR1
C3aR
pcDNA
C5aR1
C3aR
pcDNA
C5aR1
C3aR
pcDNA

**Figure S21: Surface expression of C3aR and C5aR1 in different assays.**

**A.** Surface expression of C3aR and C5aR1 in GloSensor assay (mean±SEM; n=4; normalized as fold over mock-transfection). **B.** Surface expression of C3aR and C5aR1 in βarr1 (mean±SEM; n=4) and βarr2 (mean±SEM; n=4) recruitment assay (normalized as fold over mock-transfection).
